# Sharks, Rays, & MPAs: a global assessment of marine protected area coverage in national waters across species ranges

**DOI:** 10.64898/2026.03.18.712493

**Authors:** Amanda E. Arnold, Jay H. Matsushiba, Nicholas K. Dulvy

**Affiliations:** Earth to Ocean Research Group, Department of Biological Sciences, Simon Fraser University; Burnaby, British Columbia, Canada

**Author notes:** Corresponding Author: Amanda E. Arnold, Simon Fraser University.

**Keywords:** 30x30, Area-based conservation, Biodiversity targets, Chondrichthyans, Conservation planning, Global Biodiversity Framework, Spatial prioritization, Threatened species

## Abstract

Global conservation agreements emphasize protected area coverage targets, such as the Kunming-Montreal Global Biodiversity Framework’s 30x30 target, yet their effectiveness in safeguarding biodiversity remains uncertain. We measure the intersection between marine protected area (MPAs) coverage and the distribution of sharks and rays. Using global range maps and MPA boundaries within national Exclusive Economic Zones, we calculate the percent of species’ ranges within MPAs, focusing on no-take areas. We reveal significant shortfalls in species-level protection. Within national waters, no Critically Endangered species has more than 5% of its range in no-take MPAs, and 79% of threatened species have less than 1%. We also find the WDPA contains major gaps in take-status reporting, only one third of countries (34%) report take-status of any MPAs to the WDPA, further limiting estimates of meaningful protection. These results highlight the implementation gap between global coverage targets and biodiversity outcomes, reinforcing the need for species-focused protection.

## 1. Introduction

Marine Protected Areas (MPAs) are among the primary tools for ocean conservation. The Kunming–Montreal Global Biodiversity Framework (GBF), ratified in 2022 by 196 countries, commits to protecting 30% of coastal and marine areas by 2030 (CBD 2022a). Importantly, the GBF emphasizes prioritizing areas of high biodiversity value, with guidelines highlighting the need to safeguard threatened species in order to halt and reverse biodiversity loss (CBD 2022b). However, the 30x30 target itself does not have any direct means to measure biodiversity protection, and international agreements such as the GBF can lead to the designation of MPAs in politically or economically convenient locations rather than in areas critical for biodiversity (Baldi et al. 2017; Devillers et al. 2015). This raises a key question: how well does global MPA coverage reflect the proportion of species’ ranges that are protected, particularly for species threatened with extinction?

A challenge in evaluating the suitability of MPA coverage for marine species is determining thresholds for minimum coverage. At least 10% spatial protection is necessary to deliver measurable benefits, with an average 20–40% area closure yielding maximum benefits to fisheries in previous studies (Gell and Roberts 2003; Klein et al. 2015). These thresholds align with the 30x30 framework, provided MPAs are strategically distributed and species ranges are incorporated into planning. Yet, previous studies found that 97.4% of marine species have less than 10% of their ranges covered by MPAs with restrictions on human activities (IUCN category I–IV) (Klein et al. 2015). At present, only 9.61% of the global ocean and 22.54% of national waters are under MPA protection (Protected Planet 2025), meaning countries will need to significantly expand MPA coverage to meet international biodiversity targets by 2030. In principle, average species level MPA coverage should be similar to global protection levels if MPAs are distributed strategically with the intention of protecting biodiversity.

Chondrichthyans (hereafter ‘sharks and rays’) are one of the most threatened vertebrate groups, over one-third are at risk of extinction which makes them the most threatened marine group yet evaluated (Dulvy et al. 2021). Sharks and rays threatened primarily by overfishing and bycatch (Dulvy et al. 2021). Sharks and rays are specifically sensitive due to their slow life histories (increased longevity, late maturation, and low fecundity) (Gravel et al. 2024). Using the first global assessment of sharks and rays (2014), Davidson and Dulvy (2017) found that only 12 threatened endemics had more than 10% of their species range within no-take MPAs. The 2021 global reassessment of sharks and rays resolved major geographic range uncertainties, reducing median range size from about 450,000 km² to 125,000 km² and updating species range maps and Red List statuses, thereby providing a more accurate foundation for assessing how these species are currently represented within MPAs (Dulvy et al. 2021, Dulvy et al. 2026). The combination of high extinction risk, new Red List data, and the rapid expansion of MPAs under the GBF makes this the ideal time to re-evaluate how well sharks and rays are represented in existing protected areas.

We investigate how one of the most threatened groups, sharks and rays, are encompassed by MPAs within global national waters. Specifically, we quantify the current protected area coverage of 1,158 shark and ray species by overlaying known species ranges from the IUCN Red List with current MPAs listed in the World Database of Protected Areas (WDPA). We examine the different levels of coverage based on IUCN status, paying particular attention to no-take MPA coverage of threatened species. Our aim is to provide policy makers with the current status of the spatial protection of Chondrichthyans and evidence for the need to consider species ranges in the creation of MPAs.

## 2. Methods

### 2.1 Marine Protected Area Data

The World Database on Protected Areas (WDPA) is the global repository used to assess progress toward international conservation targets (UNEP-WCMC, 2019). The database is largely self-reported, with predominantly governments and non-governmental organizations submitting protected area records (UNEP-WCMC, 2019). We primarily use the WDPA spatial polygons and two associated attributes: STATUS, which indicates the legal designation and implementation stage of each protected area, and NO_TAKE, which classifies areas based on the extent to which extractive activities are prohibited (Table S1A). The data from the WDPA used for this study was retrieved February of 2026 and details on data preparation ocan be found in Supplementary Materials Appendix A.

### 2.2 Shark and Ray Data

We used the IUCN Red List geographic range polygons for 1,158 species of sharks, rays, and chimaeras out of the 1,195 species with available shapefiles. Thirty-six species were excluded because they are found in the high seas (e.g. *Squalus boretzi*) or are primarily freshwater (e.g. *Potamotrygon falkneri*; Table S2). Only polygons where the species is currently extant were used for this analysis (Appendix A).

### 2.3 Geospatial Data Analysis

MPAs were classified into four no-take categories defined by the WDPA ‘NO_TAKE’ attribute (no-take (All), part no-take (Part), take permitted (None), and unknown (NotReported)) so that overlap area could be calculated for each category independently (Appendix A, Table S1B). To assess protection in national waters, we intersected each take-status layer with Exclusive Economic Zones (EEZs) from the Flanders Marine Institute EEZ dataset (v12).

The WDPA shapefiles were loaded into a DuckDB v1.4.2 database and most operations were conducted with the DuckDB Spatial Functions. Additional analyses were conducted with R (v4.4.2) using the sf (v1.0.20) and terra (v1.8.60) packages.

For each of the 1,172 species, range polygons were spatially clipped to the boundaries of the EEZs in which the species occurs (range within national waters). We then selected all WDPA MPAs falling within the species’ EEZ-clipped range polygon. For every species–EEZ pair, we calculated the area of overlap between the clipped species range and each MPA category. These four overlap areas were then compared with the total area of the species’ range in that EEZ to derive the percentage of protection within national waters. These spatially standardized outputs formed the basis for calculating the percentage of each species’ range protected within national waters globally. To obtain global values for each species, we summed the clipped ranges and area of overlap for each of the four MPA categories across all EEZs (yielding four global MPA coverage values per species). All areas were calculated using an ellipsoidal model of the earth. Full code and package details are provided in the Supplementary Materials.

## 3. Results

### 3.1 What is the global MPA coverage of sharks and rays in national waters?

We find that only six threatened species have more than 10% of their range within national waters covered by no-take marine protected areas (MPAs) (*Zearaja maugeana,* EN; *Centrophorus tessellatus,* EN; *Centrophorus harrissoni*, EN; *Bathyraja irrasa*, VU; *Brachaelurus colcloughi*, VU; *Squatina albipunctata*, VU; Figure 1A, Table 1). Most species across all threat categories have less than 1% of their range covered by no-take marine protected areas (Figure 1A). Of Critically Endangered (CR) sharks, 90.6% of species have less than 1% of their species range covered by a no-take MPA (Figure 2). Similarly, 88.5 % of CR rays have less than 1% of their species range protected by no-take MPAs. When we examine all threatened species, 79% have less than 1% of their range within national waters covered by no-take MPAs. When we consider all MPAs reported to the WDPA the story changes (Figure 1C). The average species overlap increases to a median of 8.6% and mean of 15%. While we would expect this to be closer to the current global coverage of 22.54% in national waters, this is still much higher than the no-take coverage of species (Figure 1A) (Protected Planet 2025). However, across all IUCN threat categories most species have less than 10% of their range covered (Table 1). We classified species with less than 1% of their geographic range occurring within MPAs as gap species, as protection at this level is unlikely to provide meaningful conservation benefit. Using this threshold, 78% of species were classified as no-take gap species (Table 1). A full list of gap species is provided in SM Appendix D.

**Figure 1:**
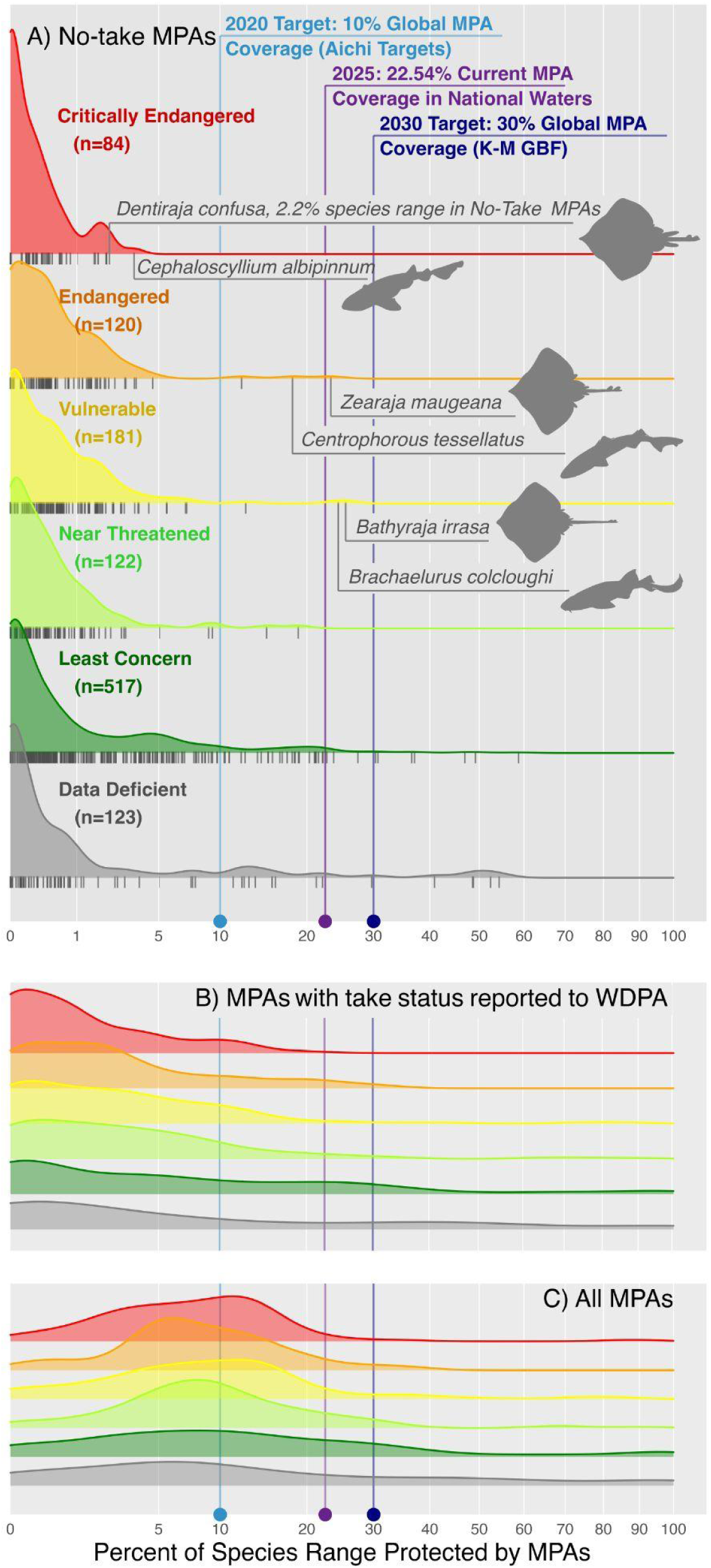
MPA coverage of shark and ray species ranges. A) Species range coverage by No-Take MPAs, separated by species’ IUCN status, selected silhouettes represent the shark and ray species with the highest MPA coverage in each threatened IUCN category (Critically Endangered, Endangered, and Vulnerable). Reference lines have been added to illustrate MPA coverage (purple) and various goals for coverage (blues). B) shows MPAs that ate labeled as No-Take, Part No-Take, and Take in the WDPA database C) shows all MPAs in the WDPA database including those where take status is not reported.

**Figure 2:**
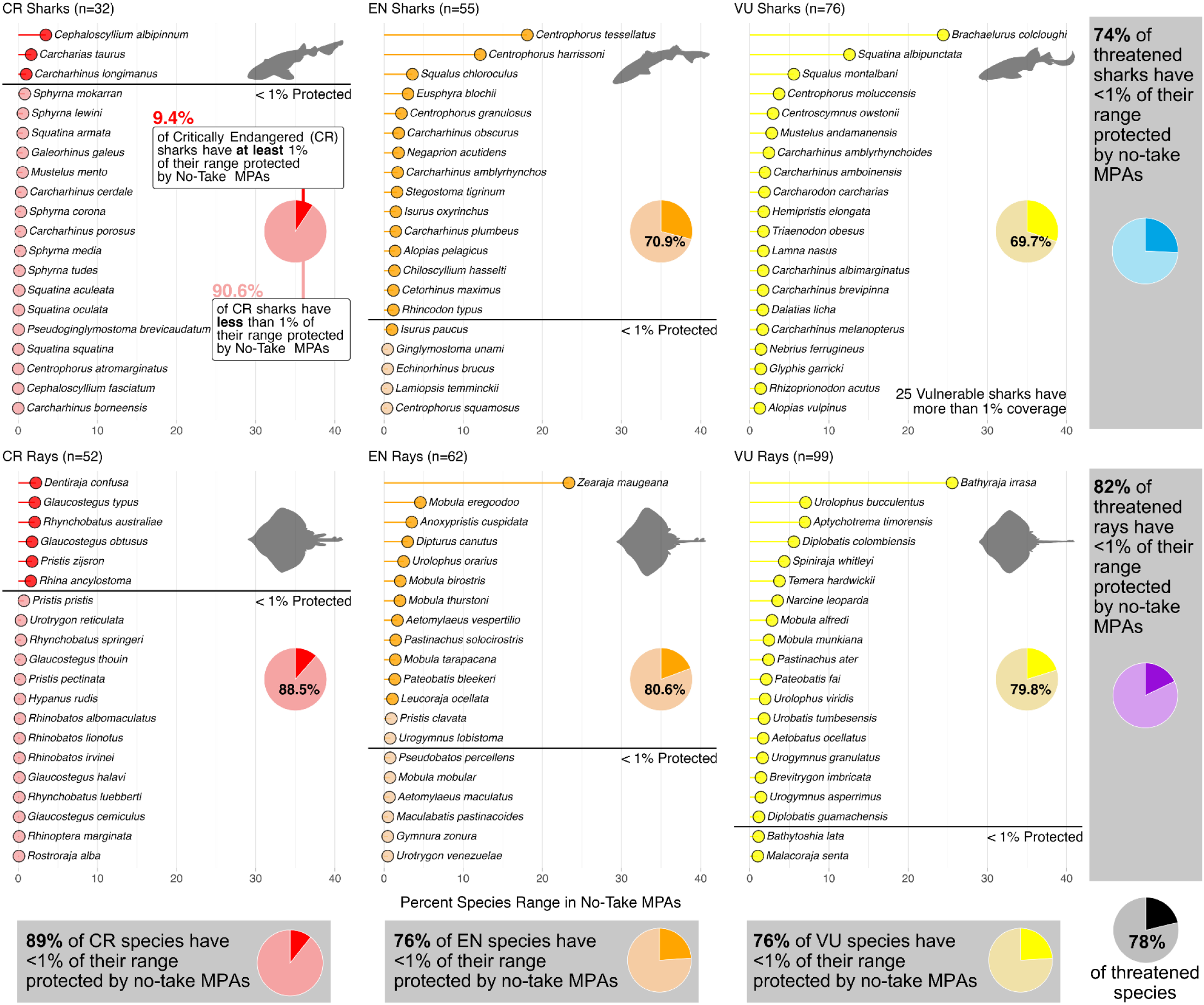
Shark and ray species with highest No-Take MPA coverage. For threatened sharks and rays (IUCN Critically Endangered, Endangered, and Vulnerable) the 20 species with the highest percentage of their species range covered by No-Take MPAs are depicted on the Y-axis. The black horizontal lines on each chart show the cut off where species below this line have less than one percent of their species range covered by No-take MPAs. Pie charts show the percentage of species in each category that have less than one percent of their range covered by No-Take MPAs. Note that the Vulnerable sharks category has more species over 1 percent than are depicted on this graph.

**Table 1:** Percentage of shark and ray species in each IUCN Red List category with less than 10% of their range protected within no-take and all MPAs and percentage of no-take gap species.

| IUCN Threat Category | Percent of species with less than 10% of range in no-take MPAs | Percent of species with less than 10% of range in any MPA | Percent of species that are no-take gap species (<1% in no-take MPAs) |
| --- | --- | --- | --- |
| CR | 100% | 60.7% | 89.3% |
| EN | 97.4% | 63.2% | 76.0% |
| VU | 98.3% | 57.5% | 76.0% |
| NT | 98.4% | 59.0% | 82.8% |
| LC | 92.8% | 51.1% | 72.7% |
| DD | 87.5% | 54.2% | 77.5% |

### 3.2 What is the extent of “Take-status” reporting in the WDPA?

Most species coverage comes from MPAs that do not report their take-status to the WDPA. Nearly half of the global MPA estate (47.8% by area) is classified as ‘Not Reported.’ Among countries with MPAs in their EEZs, only about one-third (34%) provide take-status information for any of their MPAs (Figure S3). When we account for overlapping MPA shapefiles within the WDPA dataset, the percentage of area classified as ‘Not Reported’ decreases but remains substantial (34.3%). Coverage information therefore varies across species. For some species, all MPA overlap falls within one of the three reported categories (no-take, part no-take, or take permitted), while for others, all coverage occurs within MPAs lacking take-status reporting (Figure 3).

**Figure 3.**
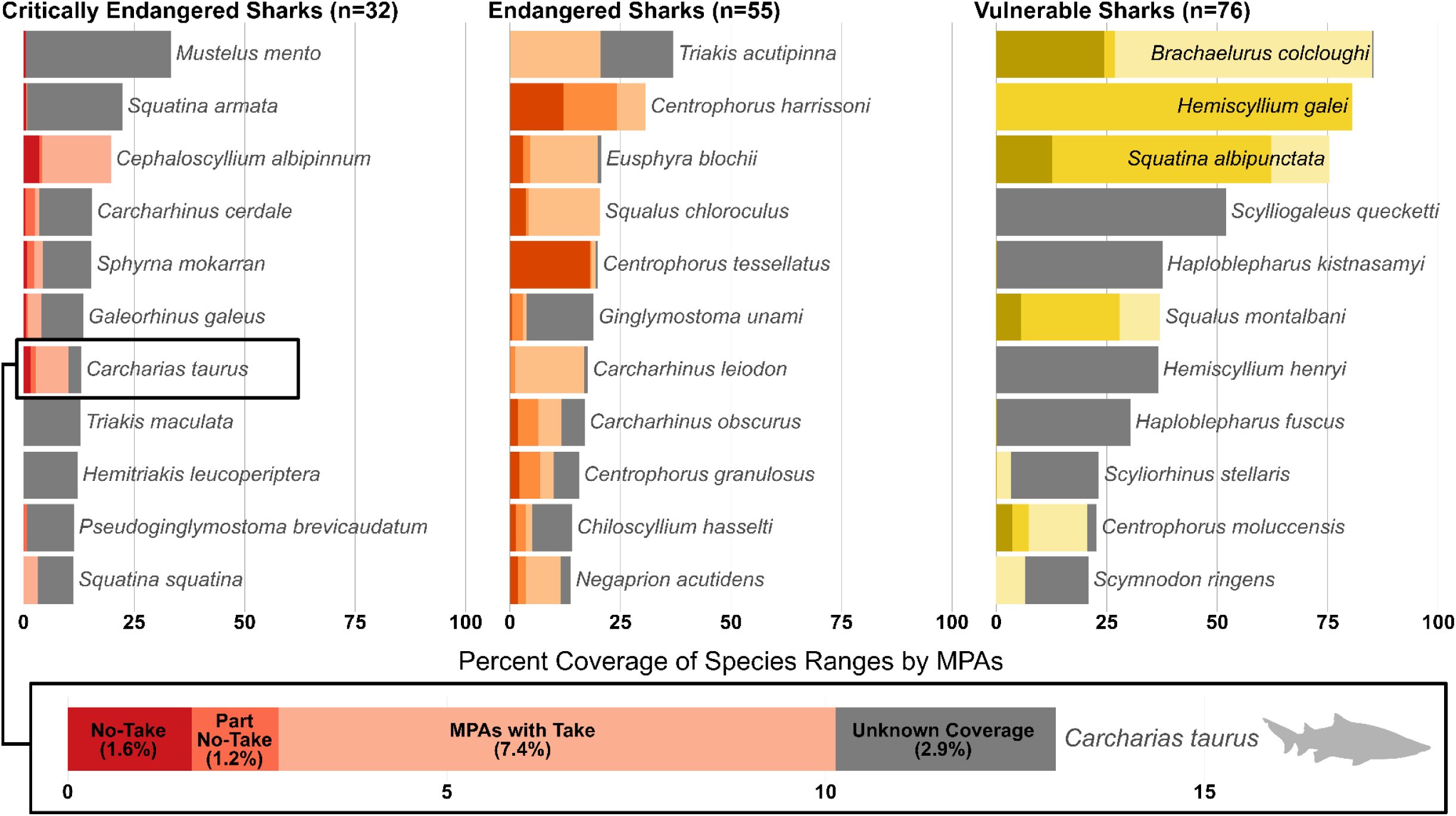
Total MPA coverage of the eleven shark species with the highest overall protected-area overlap within each threatened IUCN category. Bars show the percent of each species’ range within national waters covered by MPAs, separated by take-status classes (No-Take, Part No-Take, Take Permitted, and Not Reported). Species are ordered from highest to lowest total MPA coverage within each threat group (Critically Endangered, Endangered, Vulnerable).

### 3.3 How well is my study species protected? – SharkRayMPAexplorer

Because of the way we calculated global coverage within national waters, we also have these results at smaller geographical units (i.e., regions, subregions, and countries) and can group them by higher taxonomic levels (i.e., subclass, order, family, genus). This data are available via a RShiny app (found at https://amandaarnold.shinyapps.io/SharkRayMPAexplorer). With this tool, users are able to answer a wide breadth of questions relevant to researchers and policymakers. For example, what is the <u>mean</u> coverage of no-take MPAs for threatened species in New Zealand? (4.03%) or what is the <u>median</u> no-take percent coverage for Rhinobatidae within global national waters? (0.01%). Examples of these queries can be found in Supplementary Materials (Figure S1 & S2).

## 4. Discussion

Sharks and rays remain poorly represented within existing MPAs, even as global MPAs continue to expand under the GBF’s biodiversity targets–demonstrating that current global MPA coverage does not result in meaningful protection for most threatened species (78% under 1% covered; Figure 2). However, there is a large gap in our understanding of this coverage, in that most countries do not report take-status to the WDPA (Figure S3). This combination of low coverage and a lack of information is particularly concerning as we look for new areas of MPA expansion.

### 4.1 No-take MPAs

We focus on no-take MPAs because they provide the highest likelihood of practically reducing mortality for sharks and rays. Partial-take or multi-use MPAs often fall short of delivering meaningful protection for a variety of reasons. Even in regions where shark fishing is prohibited, the incidental capture of sharks and rays remains a major conservation concern. Bans on shark fishing do not necessarily prevent fishing pressure from altering shark populations, as bycatch can still occur at levels that affect population structure (Shea et al. 2023; Vianna et al. 2016). A second concern is the physiological sensitivity of species with high post-release mortality; management measures need to limit the frequency of fisheries interactions in order to avoid the risk of incidental catch. Hammerheads (Sphyrnidae) and devil rays (Mobulidae) have exceptionally high post-release mortality yet, on average, only 0.55% and 2.3% of their range within national waters falls inside no-take MPAs respectively (Ellis et al. 2017; IATTC 2023). Post-release mortality is also high in deepwater sharks, which are subject to high physiological stress in their landing (Finucci 2024). Because species with low intrinsic rates of population increase are highly sensitive to additional mortality, even modest levels of bycatch or post-release mortality can contribute to population declines (Gilman et al. 2022). These examples illustrate why no-take MPAs are an essential tool for reducing population-level impacts. However, the extremely low no-take coverage we report across most threatened species suggests that current MPA networks are not positioned to deliver the intended conservation benefits for most sharks and rays.

### 4.2 Lessons from example species

One threatened species that stands out in our analysis of individual MPA coverage is *Zearaja maugeana* (Maugean skate) (Grant et al. 2025). This species has a high proportion of its range within no-take MPAs (21.25%) when compared to other threatened species in our dataset. Endemic to Tasmania, Australia, the Maugean Skate has an extremely limited distribution and has only been recorded in a single estuarine system (Macquarie Harbour) over the past three decades. However, this example highlights the limits of MPA coverage alone. While the species is occasionally caught in commercial gillnet fisheries, it is prohibited from retention and post-release mortality is generally low (Grant et al. 2025). Instead, the Maugean Skate’s primary threat stems from environmental degradation caused by industrial activities within Macquarie Harbour (Lyle et al. 2014). This includes declining water quality and hypoxia driven by aquaculture and other localized stressors (Lyle et al. 2014). In such cases, conservation efforts must extend beyond spatial protection to include comprehensive environmental regulations.

The Critically Endangered whitefin swellshark (*Cephaloscyllium albipinnum*), another threatened Australian endemic, further illustrates this point. While it has the highest no-take MPA coverage for a Critically Endangered species (3.5%; Figure 1A), the spatial protection appears to be fragmented and incidental (Figure 4). That is, the MPAs were not designed for this species’ protection but happen to overlap with its limited range. This raises concerns about the effectiveness of such protections, underscoring the importance of integrated management strategies rather than reliance on MPAs alone. It is critical to match conservation tools to both the life history traits and threat profiles of individual species. Where detailed ecological and threat information exists, it should inform whether MPAs, fisheries regulations, environmental controls, or a combination thereof are most appropriate.

**Figure 4.**
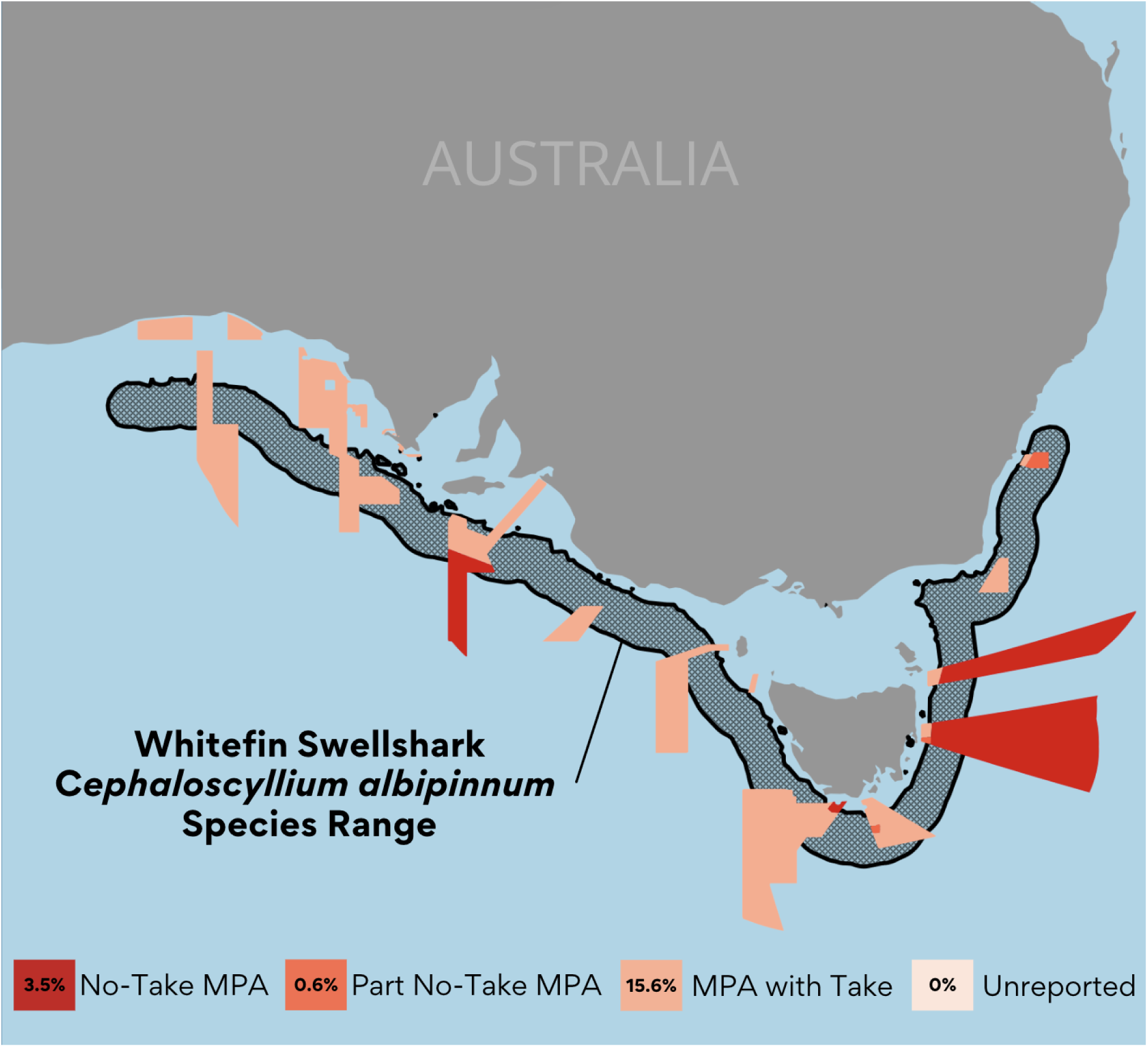
Species range and overlapping marine protected areas of the Whitefin Swellshark (*Cephaloscyllium albipinnum*) by MPA take-status category. The map shows the species’ range and all intersecting MPAs classified as No-Take, Part No-Take, Take Permitted, or Not Reported in the WDPA.

### 4.3 MPA coverage for shark conservation

The suitability of MPAs as a conservation tool is often inversely related to range size. Range-restricted species are more likely to benefit from suitable MPAs, where wide-ranging species are less likely to realize protection from spatial measures alone (Dulvy et al. 2026, Goetze et al. 2024). We use species range to estimate coverage because range polygons remain the only globally consistent spatial dataset available for all sharks and rays, enabling comprehensive cross-taxa comparisons. Additionally, of the 1,158 species we examine in this study, only around 115 have any acoustic or satellite tracking studies (Matley et al. 2024).

Although range size may overestimate true area of occupancy for some species, it provides the most comparable and scalable metric currently available for global assessment. While we reference a 10% coverage metric, it should be treated as a general benchmark rather than a universally biologically meaningful threshold. Uniform targets overlook differences in extinction risk, mobility, and geographic restriction. Faure-Beaulieu et al. (2023), suggests a range of 30% to 60% coverage with the higher end reserved for CR endemics. Additionally, the implementation of complementary conservation strategies, such as fisheries management measures, increases the benefits of fully-protected areas (Goetze et al. 2024).

### 4.4 Policy Recommendations

We have found that within global national waters no Critically Endangered shark or ray has more than 5% of its range covered by no-take MPAs and, of threatened species, 79% have less than 1% of their range covered. Of the countries who submit MPAs to the WDPA, only 34% report any take-status to the WDPA. As we move toward the 30x30 target under the GBF, these results highlight areas where policy changes could greatly improve biodiversity outcomes:

1. *Monitoring for biodiversity targets in the GBF.* Despite emphasizing the need to prioritize areas of particular importance for biodiversity in Targets 1 and 3, the GBF currently includes no indicators for biodiversity outcomes, and thus no way to connect increased protection of habitat to improvements in species status in its Monitoring Framework (CBD, 2022b). While global area-based targets lay a critical foundation, their effectiveness depends on integrating clear targets for species-level protection. The ability to report MPA coverage to a species level, creates practical information on our progress towards biodiversity targets. This research provides a replicable framework for evaluating species’ spatial coverage and can help guide biologically informed expansion of protections based on species range.
2. *Requiring MPA submissions to include take-status when reporting to the WDPA.* Standardizing take-status reporting would greatly improve the global ability to assess the functional protection that MPAs provide. Requiring take-status as part of national submissions would increase the transparency of global MPA reporting and allow for more accurate evaluations of biodiversity targets, particularly for species at risk of extinction.
3. *Promoting biodiversity-driven decision making when delineating MPAs.* Strengthening MPAs requires biodiversity to be accounted for in the initial design stages. Aligning MPAs with species distributions and Key Biodiversity Areas could be methods to ensure new protections deliver biodiversity benefits for threatened species (KBA Secretariat 2024). Systematic conservation planning can also be used to identify areas where species lack sufficient protection, set species specific protection targets based on threatened status, and highlight critical areas where targeted spatial protection would yield the greatest conservation benefit (Faure-Beaulieu et al. 2023). For sharks and rays specifically, there is an opportunity to leverage Important Shark and Ray Areas to create more tailored protections for these threatened and geographically restricted species (Boyd et al. 2025; Hyde et al. 2022).

### 4.5 Future Research

We encourage future research to compare the MPAs included in the WDPA with those found in other global databases, such as the Marine Protection Atlas (MPAtlas) (Pike et al. 2024, Protected Planet 2024). While our objective here was to assess species-level coverage using the WDPA, the dataset currently used to set global conservation benchmarks, evaluating species coverage using databases with independently assessed MPA classification schemes would provide valuable additional insight and improve our understanding of areas where take-status is not reported. While the MPAtlas dataset does not include all MPAs (as we do here with the WDPA database) every entry has a protection level reported.

## Acknowledgements and Data

This research was supported by the Natural Sciences and Engineering Research Council of Canada, the Weston Family Foundation, and the Abbott-Fretwell Graduate Fellowship in Fisheries Biology. We declare no financial or personal conflicts of interest that could influence the objectivity of this work. All data and code will be made publicly available in a GitHub repository upon publication. Species range data were obtained from the IUCN Red List and MPA boundaries from the WDPA (IUCN 2025; UNEP-WCMC and IUCN 2025).

