## Supplementary Materials for "Sharks, Rays, & MPAs: a global assessment of marine protected area coverage in national waters across species ranges"

**Contents:**

**Appendix A:** Data cleaning process for WDPA and IUCN Redlist data
**Table S1:** Attributes from the World Database of Protected Areas (WDPA) used in this analysis.
**Table S2:** Species excluded from analysis due to their ranges and reasoning behind exclusion.
**Appendix B:** Additional results
**Appendix C:** Data and Code Accessibility
**Figure S1:** Screen capture of example query from SharkRayMPAexplorer Shiny App: What is the MPA coverage of each no-take category for threatened species in New Zealand?
**Figure S2:** Screen capture of example query from SharkRayMPAexplorer Shiny App: What is the average no-take percent coverage for Rhinobatidae within global national waters?
**Figure S3:** Map of countries with any MPAs in the WDPA with take status.
**Appendix D**: List of gap species.

### Appendix A: Data cleaning process for WDPA and IUCN Redlist data

#### *WDPA Data Cleaning*

For the purposes of this study we are only examining marine protected areas (MPAs) from the WDPA database; OECMs were not included as part of this analysis. We filtered the WDPA database to 'Marine' and 'Coastal' categories of protected areas, excluding 'Terrestrial'. We excluded any MPAs that were not yet active, as 'proposed' MPAs are also in the WDPA data set (Table S1B). Additionally, we only included MPAs within exclusive economic zones (EEZs) as only approximately 1.5% of the high seas has currently implemented MPAs (MPA Guide, 2023). We also exclude UNESCO Biosphere reserves from the analysis, following best practices. There are many overlapping MPAs in the WDPA dataset, this is likely due to different jurisdictional regulations placed over an area (for example a local no-take MPA could be placed within a federal protected area). We intersected MPA polygons with one another ordinally, according to take value (no-take > part no-take > take permitted > not reported), and assume the most restrictive conditions apply to a covered area. For example, if an area has overlapping 'no-take' and 'not reported' polygons we will assume that area is no-take, resulting in a non-overlapping set of unique features. This means our analysis is highly conservative and the estimates represent an upper limit of protection.

#### *IUCN Redlist Data Cleaning*

First, we only included species range polygons from the IUCN Redlist that were classified as "Extant (resident)" where PRESENCE code is equal to 1. This excludes areas that are "Possibly Extant (resident)", "Presence Uncertain", "Possibly Extinct", and "Presence Uncertain & Origin Uncertain". As with the WDPA data, species ranges were clipped to Exclusive Economic Zones (EEZs). This step primarily affected the 40 species with primary or secondary pelagic habitat, out of the 1,172 species used in the analysis. Because this study focuses on species coverage within EEZs, we excluded range areas in the high seas to avoid inflating or distorting coverage metrics. Additionally, some species are freshwater or fall outside of EEZs entirely, these species were excluded from analysis (Table S1). All polygons used for analysis had an ORIGIN code equal to 1 and a SEASONALITY code equal to 1.

**Table S1:** Attributes from the World Database of Protected Areas (WDPA) used in this analysis. Tables adapted from UNEP-WCMC (2019).

**A)** Overview of non-spatial attributes used

| No | Field Name | Requirement | Provided by | Type | Accepted values |
|---|---|---|---|---|---|
| 1 | WDPAID | Minimum | UNEP-WCMC | Number (Double) | Assigned by UNEP-WCMC. Unique identifier for a protected Area. |
| 4 | NAME | Minimum | Data provider | Text (String) | Name of the protected area (PA) as provided by the data provider |
| 11 | MARINE | Minimum | Data provider | Text (String) | **Possible values**: 0 (100% Terrestrial PA, 1 (Coastal: marine and terrestrial PA), and 2 (100% marine PA).<br><br>**Selected values:** 1 (Coastal: marine and terrestrial PA), and 2 (100% marine PA). |
| 16 | NO_TAKE | Complete | Data provider | Text (String) | **Possible Values:** All, Part, None, Not Reported, Not Applicable (if Marine field = 0).<br><br>**Selected Values:** All, Part, None, Not Reported, Not Applicable (if Marine field = 0). |
| 18 | STATUS | Minimum | Data provider | Text (String) | **Possible Values:** Proposed, Inscribed, Adopted, Designated, Established.<br><br>**Selected Values:** Inscribed, Adopted, Designated, Established. |
| 29 | ISO3 | Minimum | Data provider | Text (String) | ISO 3166-3 character code of country or territory where the PA is located. |

**B)** Description of allowed values for the NO_TAKE attribute

| If Marine is: | Accepted Values | Description |
|---|---|---|
| 1/2 | All | For marine protected areas, No Take is listed as to whether all, part or none of the protected area is no take. (No take means that the taking of natural resources, inclusive of all methods of fishing, extraction, dumping, dredging and construction, is strictly prohibited in all or part of a marine protected area.) |
| | Part | |
| | None | |
| | Not Reported | For marine protected areas where it is not known whether there is no take, 'Not Reported' is listed. |
| 0 | Not Applicable | For non-marine protected areas 'Not Applicable' is listed. |

**C)** Description of allowed values for the STATUS attribute

| Accepted Values | Description |
|---|---|
| Proposed | Is in a process to gain recognition or dedication through legal or other effective means. It should be noted that a site may be managed as a protected area while proposed, as the national legal processes of designation may take a long time. |
| Inscribed | Only applicable for protected areas designated under the World Heritage Convention. |
| Adopted | Only applicable to protected areas designated as Specially Protected Area of Marine Importance under the Barcelona Convention. |
| Designated | Is recognized or dedicated through legal means. Implies specific binding commitment to conservation in the long term. |
| Established | Recognized or dedicated through other effective means. Implies commitment to conservation outcomes in the long term, but not necessarily with legal recognition. |

**Table S2:** Species excluded from analysis due to their ranges and reasoning behind exclusion.

| | **Species** | **Reason for Exclusion** |
|---|---|---|
| 1 | *Bathyraja arctowskii* | Only found in ABNJ |
| 2 | *Bathyraja tunae* | Only found in ABNJ |
| 3 | *Bythaelurus bachi* | Only found in ABNJ |
| 4 | *Bythaelurus naylori* | Only found in ABNJ |
| 5 | *Centroscyllium excelsum* | Only found in ABNJ |
| 6 | *Chimaera buccanigella* | Only found in ABNJ |
| 7 | *Chimaera didierae* | Only found in ABNJ |
| 8 | *Etmopterus lailae* | Only found in ABNJ |
| 9 | *Heliotrygon gomesi* | Freshwater Species |
| 10 | *Hemitrygon laosensis* | Freshwater Species |
| 11 | *Leucoraja longirostris* | Only found in ABNJ |
| 12 | *Makararaja chindwinensis* | Freshwater Species |
| 13 | *Mollisquama parini* | Only found in ABNJ |
| 14 | *Notoraja lira* | Only found in ABNJ |
| 15 | *Paratrygon parvaspina* | Freshwater Species |
| 16 | *Plesiotrygon nana* | Freshwater Species |
| 17 | *Potamotrygon adamastor* | Freshwater Species |
| 18 | *Potamotrygon albimaculata* | Freshwater Species |
| 19 | *Potamotrygon amandae* | Freshwater Species |
| 20 | *Potamotrygon amazona* | Freshwater Species |
| 21 | *Potamotrygon falkneri* | Freshwater Species |
| 22 | *Potamotrygon garmani* | Freshwater Species |
| 23 | *Potamotrygon jabuti* | Freshwater Species |
| 24 | *Potamotrygon limai* | Freshwater Species |
| 25 | *Potamotrygon pantanensis* | Freshwater Species |
| 26 | *Potamotrygon rex* | Freshwater Species |
| 27 | *Potamotrygon schroederi* | Freshwater Species |
| 28 | *Potamotrygon signata* | Freshwater Species |
| 29 | *Potamotrygon tatianae* | Freshwater Species |
| 30 | *Potamotrygon tigrina* | Freshwater Species |

| | | |
|---|---|---|
| 31 | *Potamotrygon wallacei* | Freshwater Species |
| 32 | *Rhinobatos nudidorsalis* | Only found in ABNJ |
| 33 | *Scymnodalatias oligodon* | Only found in ABNJ |
| 34 | *Squalus boretzi* | Only found in ABNJ |

**Appendix B:** Additional Results

We modelled proportional marine protected area (MPA) coverage within national waters using Bayesian zero–one inflated beta regression implemented in the `brms` package in R. Zero–one inflated beta regression was used to accommodate proportional coverage data bounded in (0,1) while explicitly modelling structural zeros and complete coverage. This allows predictor variables to influence not only average MPA coverage but also the probability of complete absence or complete coverage.

We evaluated relationships for both no-take MPA coverage (NO_TAKE = "All") and total MPA coverage (NO_TAKE = "All", "Part", "None", and "Not Reported"). Predictor variables were included, where appropriate, in the mean ($\mu$), precision ($\varphi$), and both zero- and one-inflation components of the model. This structure allowed predictors such as species range size to influence not only average proportional coverage, but also the probability of complete absence of protection or complete protection. No species exhibited complete no-take coverage (proportion = 1); therefore, for no-take models the one-inflation component was fixed (coi ~ 1), while zero inflation was estimated where applicable. Models were fit using four Markov chain Monte Carlo chains (3,000 iterations each, including 1,000 warmup iterations) with adapt_delta = 0.95 to ensure stable convergence.

Predictor variables were evaluated individually and interactions between predictors were not included. This approach allowed us to isolate the marginal relationship between each ecological or geographic predictor and MPA coverage, and results should therefore be interpreted as exploratory.

Species Range

First, we examined the relationship between total species range area (including areas in the high seas) and MPA coverage. Predicted coverage within no-take MPAs remained consistently low across the range-size gradient, indicating no strong relationship between species range area and no-take MPA coverage. In contrast, predicted total MPA coverage was highest for species with small to medium range sizes and declined for species with the largest ranges. Uncertainty was greatest for species with the smallest ranges and decreased as range size increased, indicating greater variability in coverage among range-restricted species while wide ranging species are more likely to have at least some small part of their range protected (Figure B2).

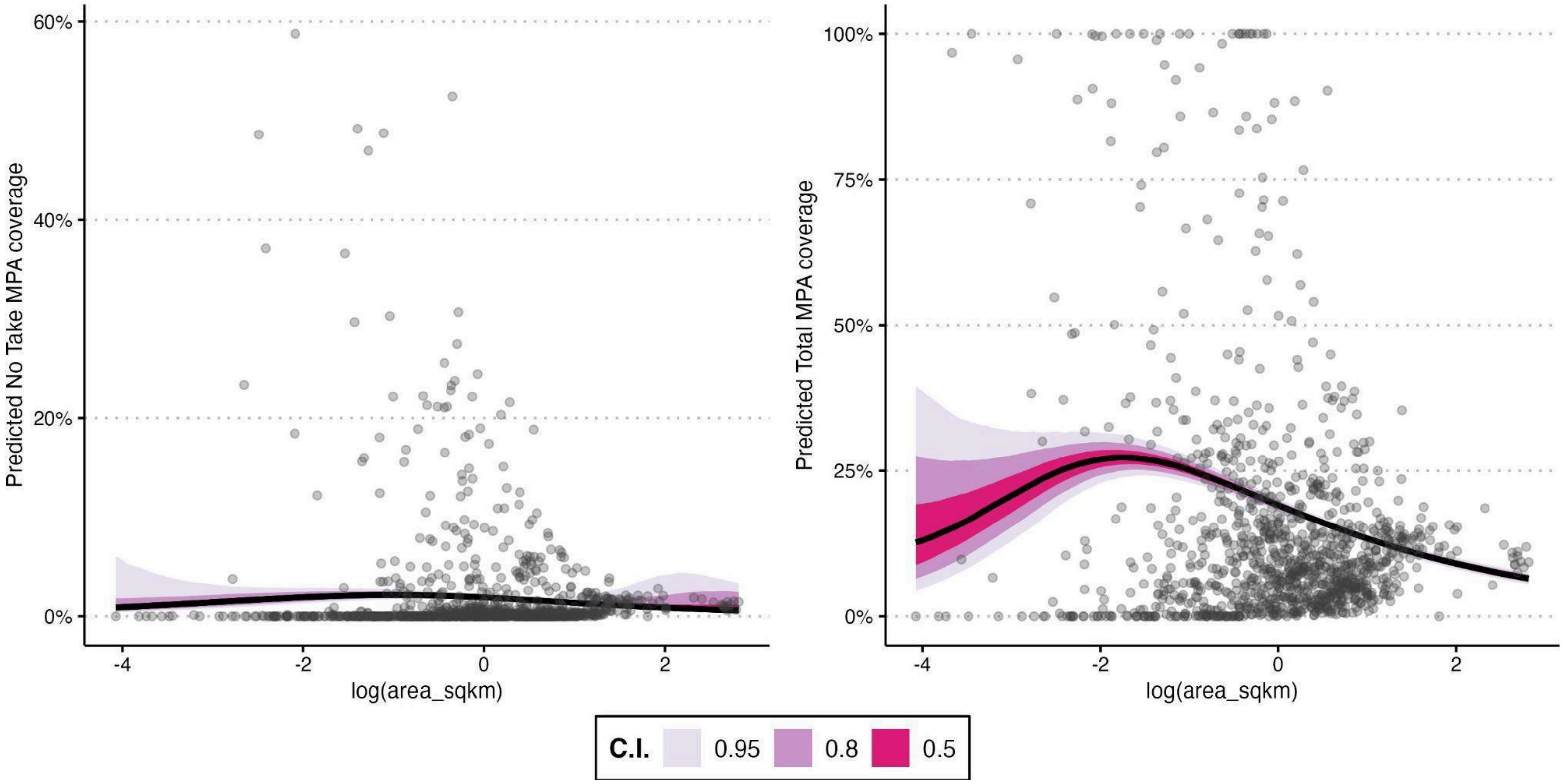


**Figure B1:** Relationship between species range size and predicted marine protected area (MPA) coverage for sharks and rays. Predictions from Bayesian zero–one inflated beta models are shown as a function of centered and scaled log-transformed median depth (m). Points represent observed species values (n = 1,158). The black line indicates the posterior median with 50%, 80%, and 95% credible intervals. The left panel shows predicted coverage within no-take MPAs only, and the right panel shows predicted coverage across all MPAs regardless of protection level.

Mid Depth

Next, we looked at the relationship between species depth midpoint (calculated as the middle point between minimum depth and maximum depth) and MPA coverage. Again, predicted coverage within no-take MPAs was consistently low across the depth gradient, indicating limited strict protection for both shallow and deep-water species. In contrast, predicted total MPA coverage increased with depth, suggesting that deeper-ranging species are more frequently included within MPAs when partially protected or multi-use areas are considered (Figure B2).

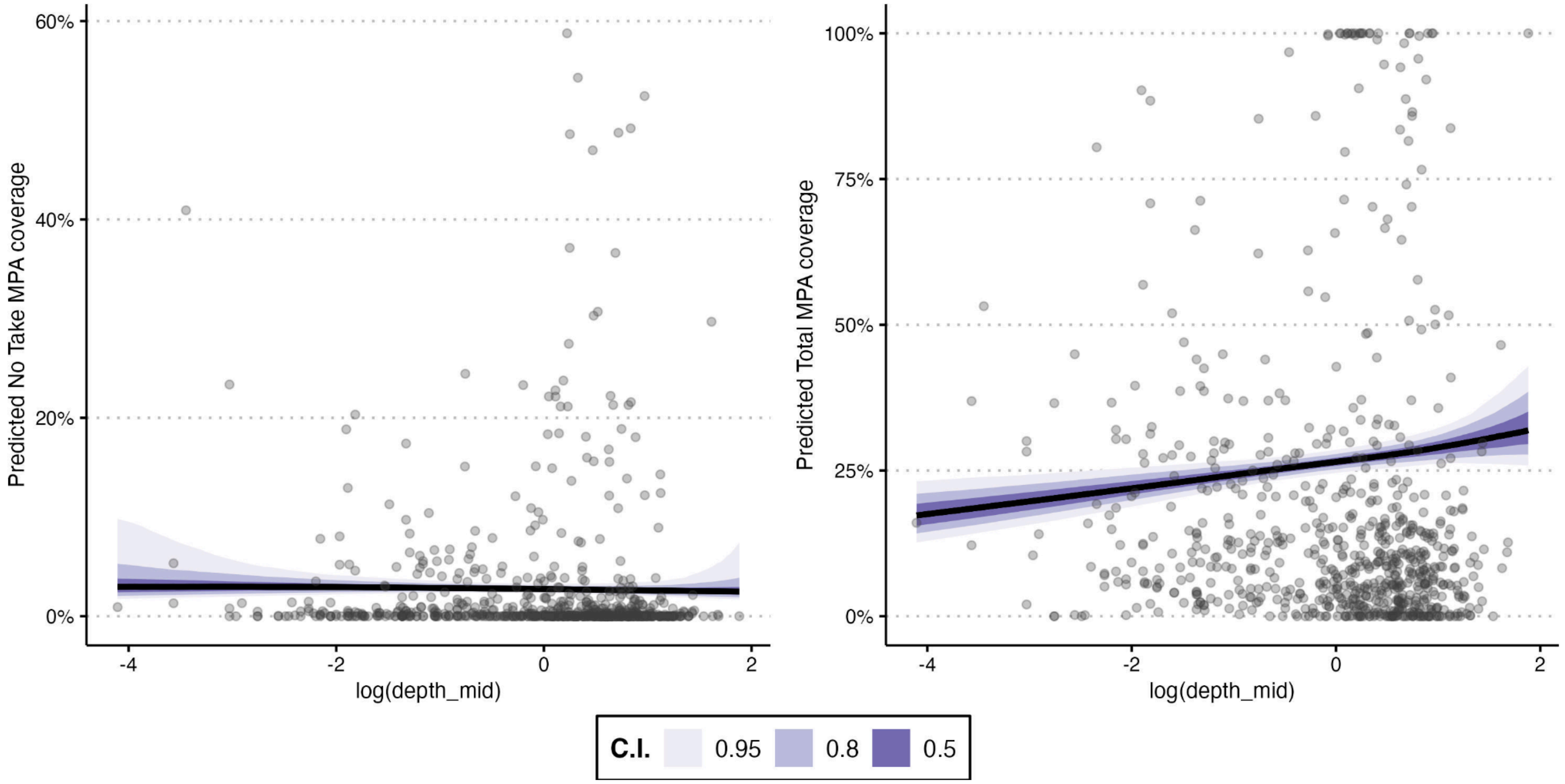


**Figure B2.** Relationship between species mid-depth and predicted marine protected area (MPA) coverage for sharks and rays. Predictions from Bayesian zero–one inflated beta models are shown as a function of centered and scaled log-transformed median depth (m). Points represent observed species values (n = 1,158). The black line indicates the posterior median with 50%, 80%, and 95% credible intervals. The left panel shows predicted coverage within no-take MPAs only, and the right panel shows predicted coverage across all MPAs regardless of protection level.

Body Class Size

Predicted coverage within no-take MPAs remained very low across all size classes, with little difference between small, medium, and large-bodied species, as defined in Dulvy et al. (2024). In contrast, predicted total MPA coverage was greatest for small-bodied species and lower for medium and large-bodied species (Figure B3).

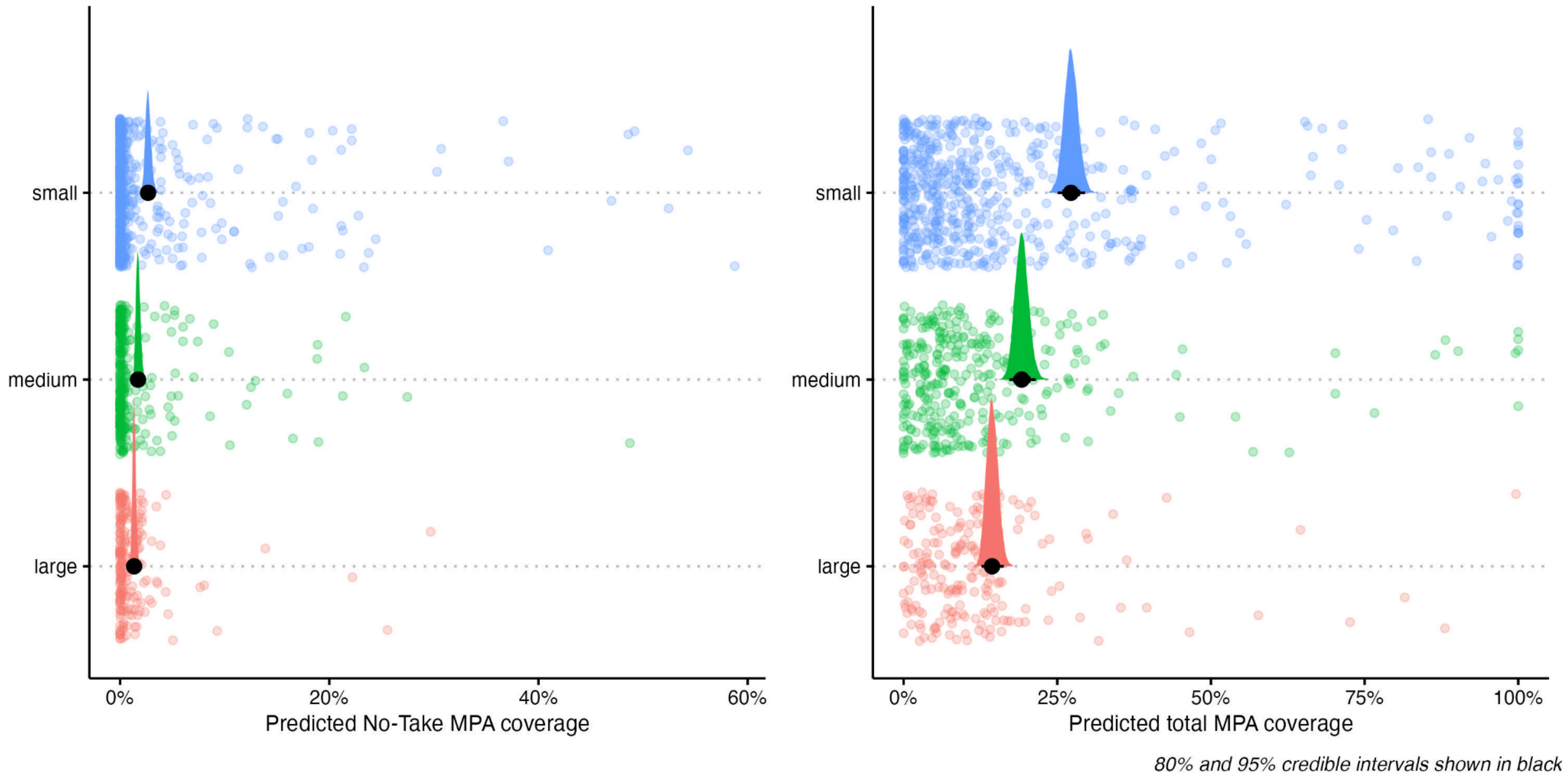


**Figure B3:** Relationship between species body size category and predicted marine protected area (MPA) coverage for sharks and rays. Predicted proportional coverage is shown for species grouped into small, medium, and large body size categories. Points represent observed species-level values (n = 1,158). Black points indicate the posterior median predicted coverage from Bayesian zero–one inflated beta models, with black intervals showing the 80% and 95% credible intervals. Colored densities illustrate the posterior distribution of predicted values for each size category. The left panel shows predicted coverage within no-take MPAs only, whereas the right panel shows predicted total MPA coverage across all MPA categories.

Primary Habitat

Predicted coverage within no-take MPAs remained very low across all primary habitats, with little difference between pelagic, deepwater, and coastal species, as defined in Dulvy et al. (2021). In contrast, predicted total MPA coverage was greatest for deepwater species and lower for pelagic and coastal species (Figure B4).

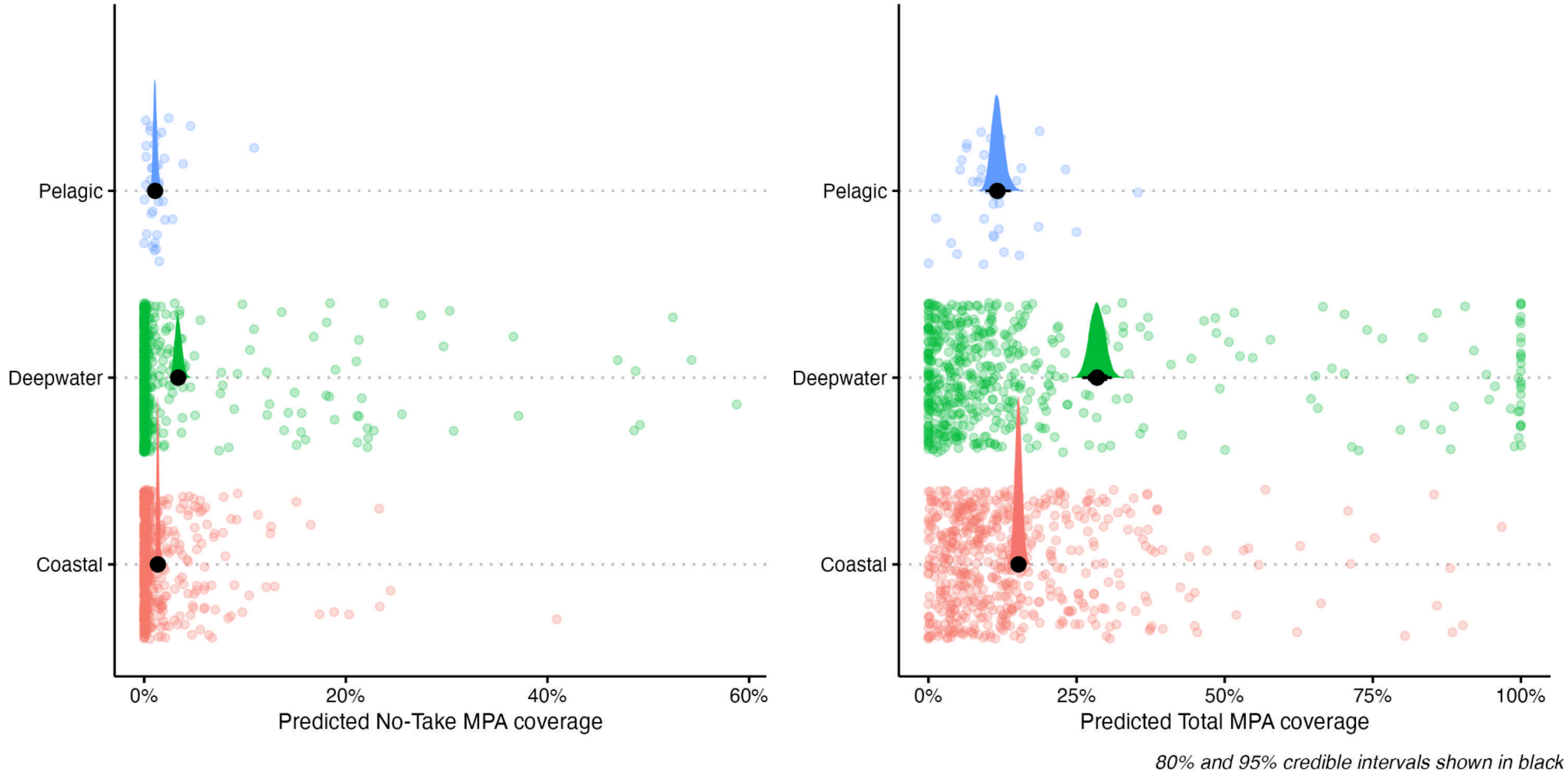


**Figure B4:** Relationship between species primary habitat and predicted marine protected area (MPA) coverage for sharks and rays. Predicted proportional coverage is shown for species grouped into pelagic, deepwater, and coastal primary habitats. Points represent observed species-level values (n = 1,158). Black points indicate the posterior median predicted coverage from Bayesian zero–one inflated beta models, with black intervals showing the 80% and 95% credible intervals. Colored densities illustrate the posterior distribution of predicted values for each size category. The left panel shows predicted coverage within no-take MPAs only, whereas the right panel shows predicted total MPA coverage across all MPA categories.

Order

We compared coverage in each taxonomic order against MPA coverage. Predicted coverage within no-take MPAs remained very low across all orders. There was also significant overlap between most orders when examining total MPA coverage (Figure B5).

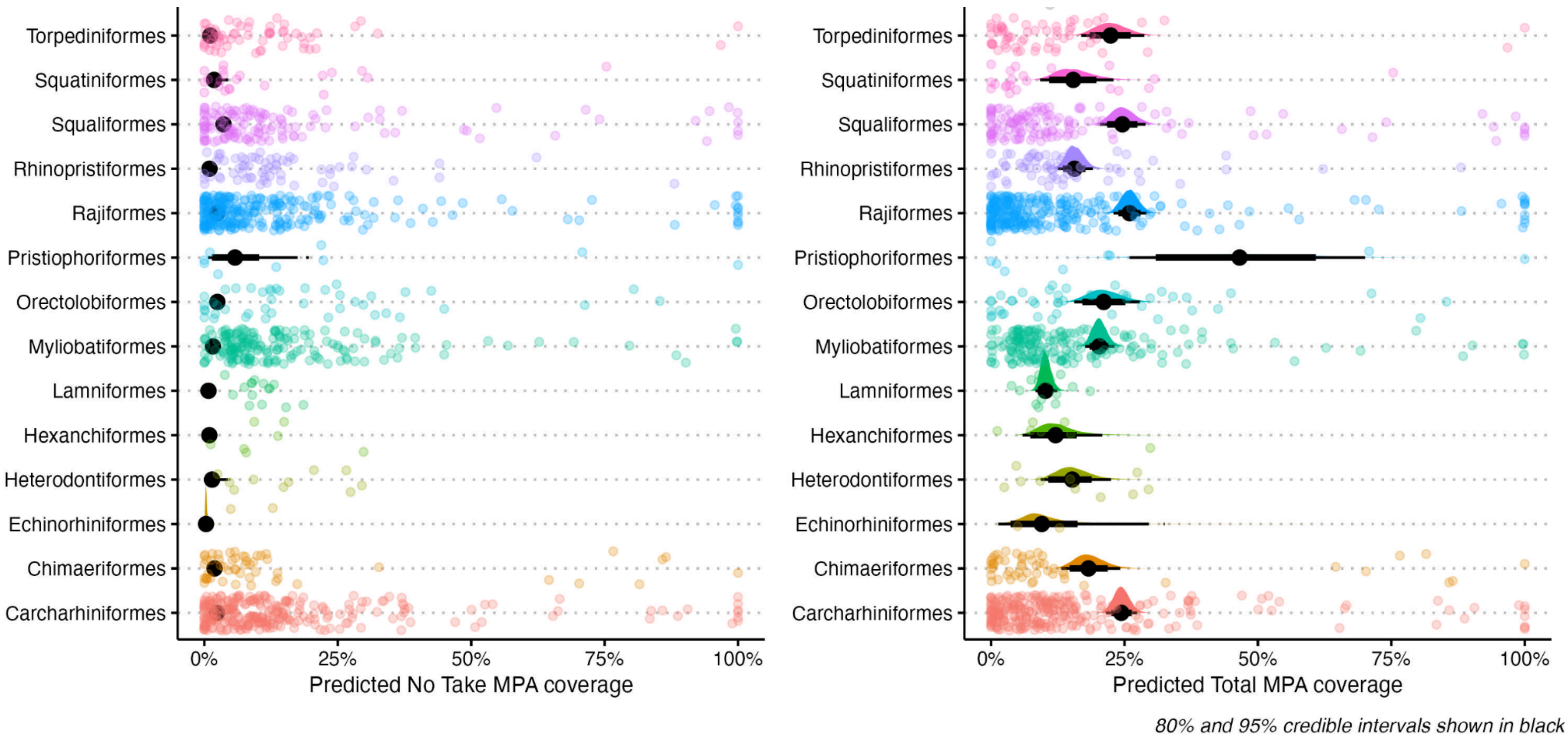


**Figure B5:** Relationship between species primary habitat and predicted marine protected area (MPA) coverage for sharks and rays. Predicted proportional coverage is shown for species grouped into pelagic, deepwater, and coastal primary habitats. Points represent observed species-level values (n = 1,158). Black points indicate the posterior median predicted coverage from Bayesian zero–one inflated beta models, with black intervals showing the 80% and 95% credible intervals. Colored densities illustrate the posterior distribution of predicted values for each size category. The left panel shows predicted coverage within no-take MPAs only, whereas the right panel shows predicted total MPA coverage across all MPA categories.

Ecomorphotype

Predicted coverage within no-take MPAs was uniformly low across most ecomorphotypes (White et al. 2022) . In contrast, predicted total MPA coverage differed more clearly among some ecomorphotypes. Several benthic-associated groups, including pristiobenthic, probenthic, and rajobenthic species, showed relatively higher predicted coverage compared to more pelagic-associated groups such as microoceanic and macrooceanic species. Despite these differences, species-level observations revealed substantial variability within most ecomorphotype categories.

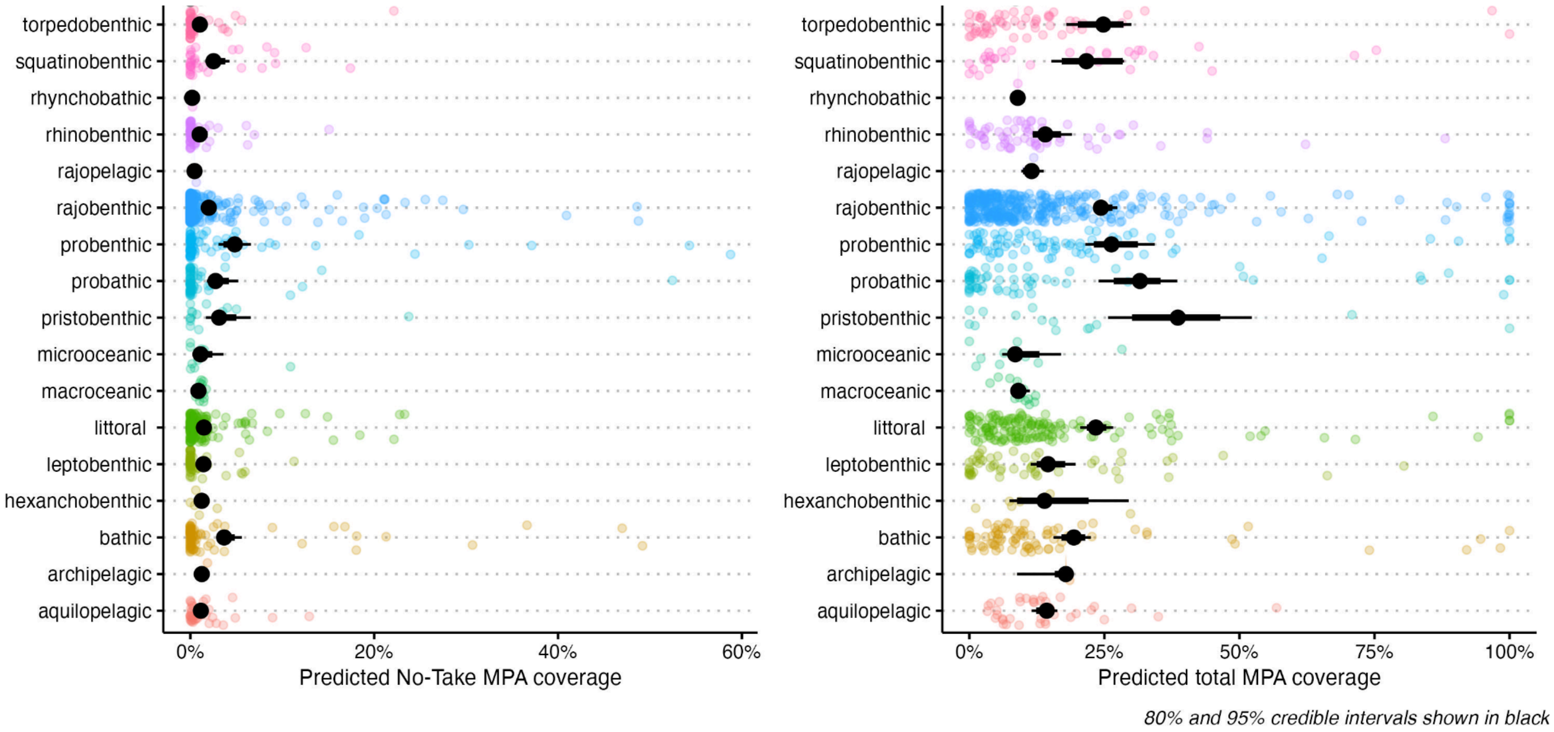


**Figure B6:** Relationship between ecomorphotype and predicted marine protected area (MPA) coverage for sharks and rays. Predicted proportional coverage is shown for species belonging to ecomorphotype categories. Points represent observed species-level values (n = 1,158). Black points indicate the posterior median predicted coverage from Bayesian zero–one inflated beta models, with black intervals showing the 80% and 95% credible intervals. Colored densities illustrate the posterior distribution of predicted values for each size category. The left panel shows predicted coverage within no-take MPAs only, whereas the right panel shows predicted total MPA coverage across all MPA categories.

<u>Endemism</u>

Predicted coverage within no-take MPAs was uniformly low when comparing endemic and non-endemic species. However, when examining total coverage endemic species have a higher predicted total MPA coverage (Figure B7).

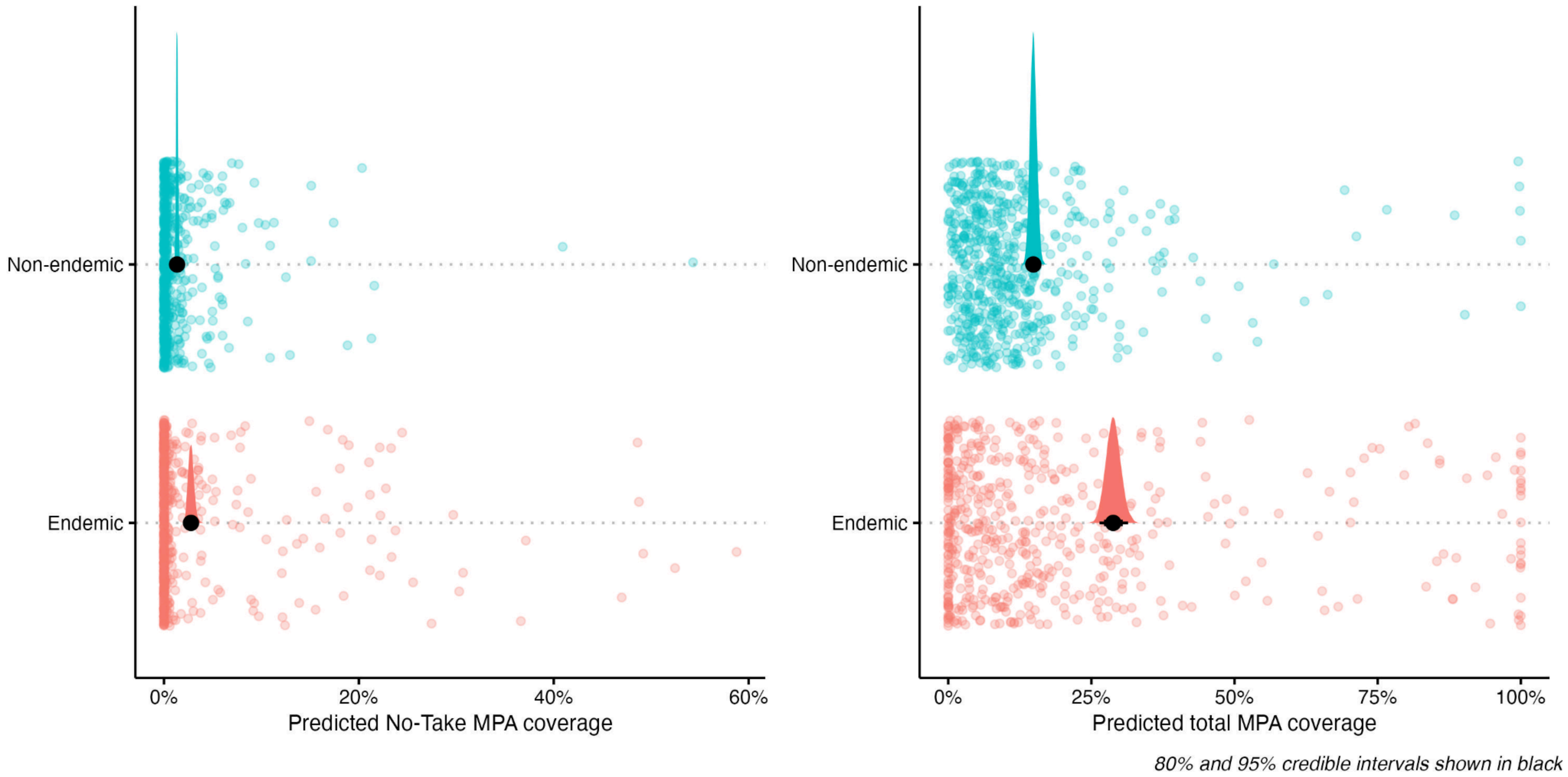


**Figure B7:** Relationship between species endemism and predicted marine protected area (MPA) coverage for sharks and rays. Predicted proportional coverage is shown for species grouped into endemic and non-endemic. Points represent observed species-level values (n = 1,158). Black points indicate the posterior median predicted coverage from Bayesian zero–one inflated beta models, with black intervals showing the 80% and 95% credible intervals. Colored densities illustrate the posterior distribution of predicted values for each size category. The left panel shows predicted coverage within no-take MPAs only, whereas the right panel shows predicted total MPA coverage across all MPA categories.

Shark Catch

For this analysis, we used reported shark catch data from the Sea Around Us Project (Dulvy et al. 2024). Average shark catch per territory was calculated from reported catch between 2010 and 2019, and mean MPA coverage was calculated for shark and ray species occurring within each territory for which catch data were available. Average shark catch per territory showed a negative relationship with predicted marine protected area (MPA) coverage. Predicted coverage within no-take MPAs was consistently low across the full range of catch values and declined slightly as average catch increased. In contrast, predicted total MPA coverage showed a stronger negative relationship with catch, with territories exhibiting higher average catch having substantially lower predicted MPA coverage. Uncertainty was greatest at lower catch values and decreased as catch increased, but the overall pattern suggests that areas with higher fishing pressure tend to have lower overall MPA coverage, consistent with residual reserves.

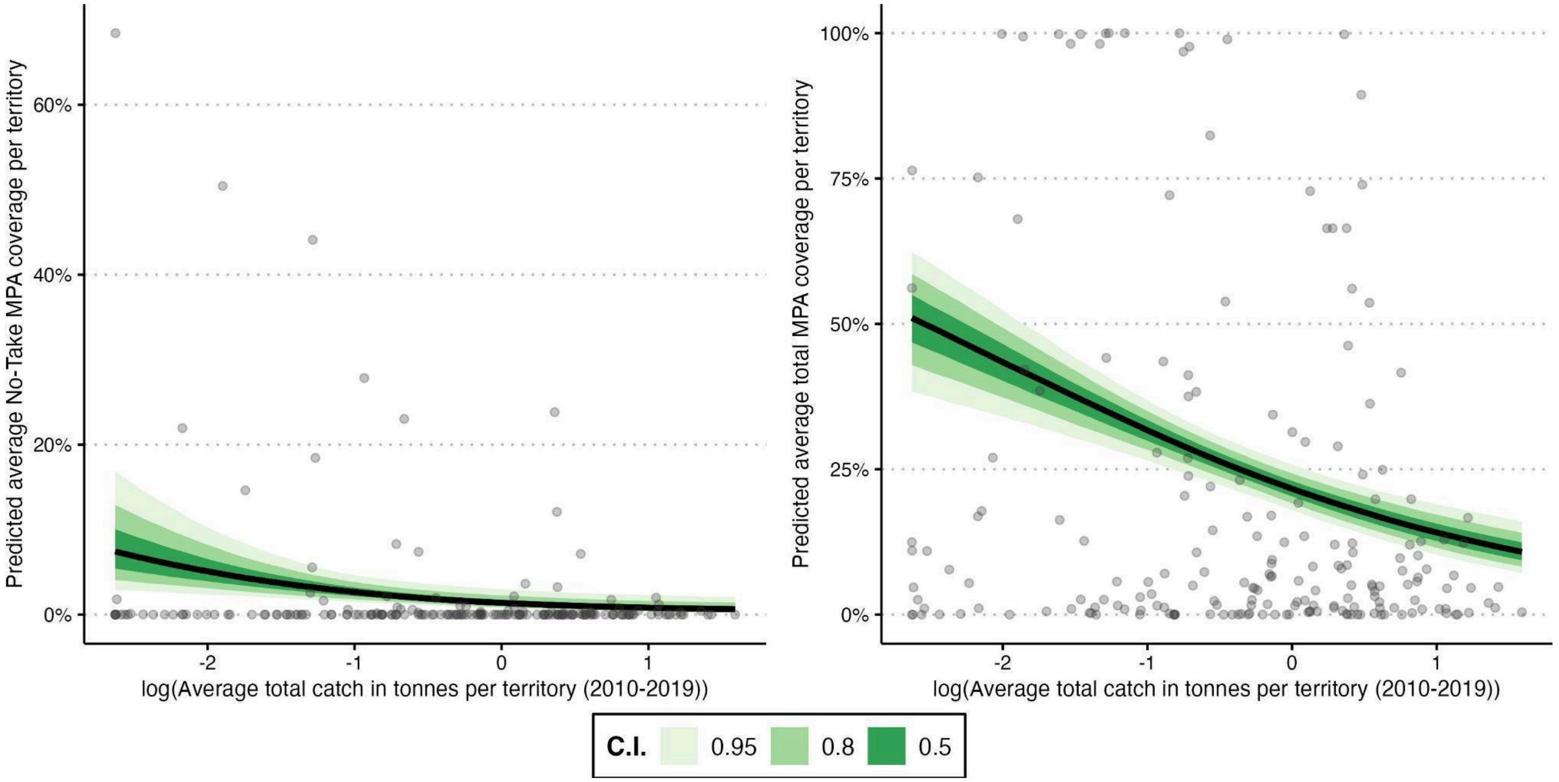


**Figure B8.** Relationship between average total shark catch and predicted average marine protected area (MPA) coverage for sharks and rays. Predictions from Bayesian zero–one inflated beta models are shown as a function of centered and scaled log-transformed average total shark catch (tonnes). Points represent mean protected area coverage of sharks and rays within a given territory (n =). The black line indicates the posterior median with 50%, 80%, and 95% credible intervals. The left panel shows predicted coverage within no-take MPAs only, and the right panel shows predicted coverage across all MPAs regardless of protection level.

Subregions

Predicted coverage within no-take MPAs was uniformly low across all subregions. In contrast, predicted total MPA coverage differed more clearly among some subregions. Several subregions, notably Western Europe, Polynesia, and Australia and New Zealand, showed higher predicted coverage compared to other subregions.

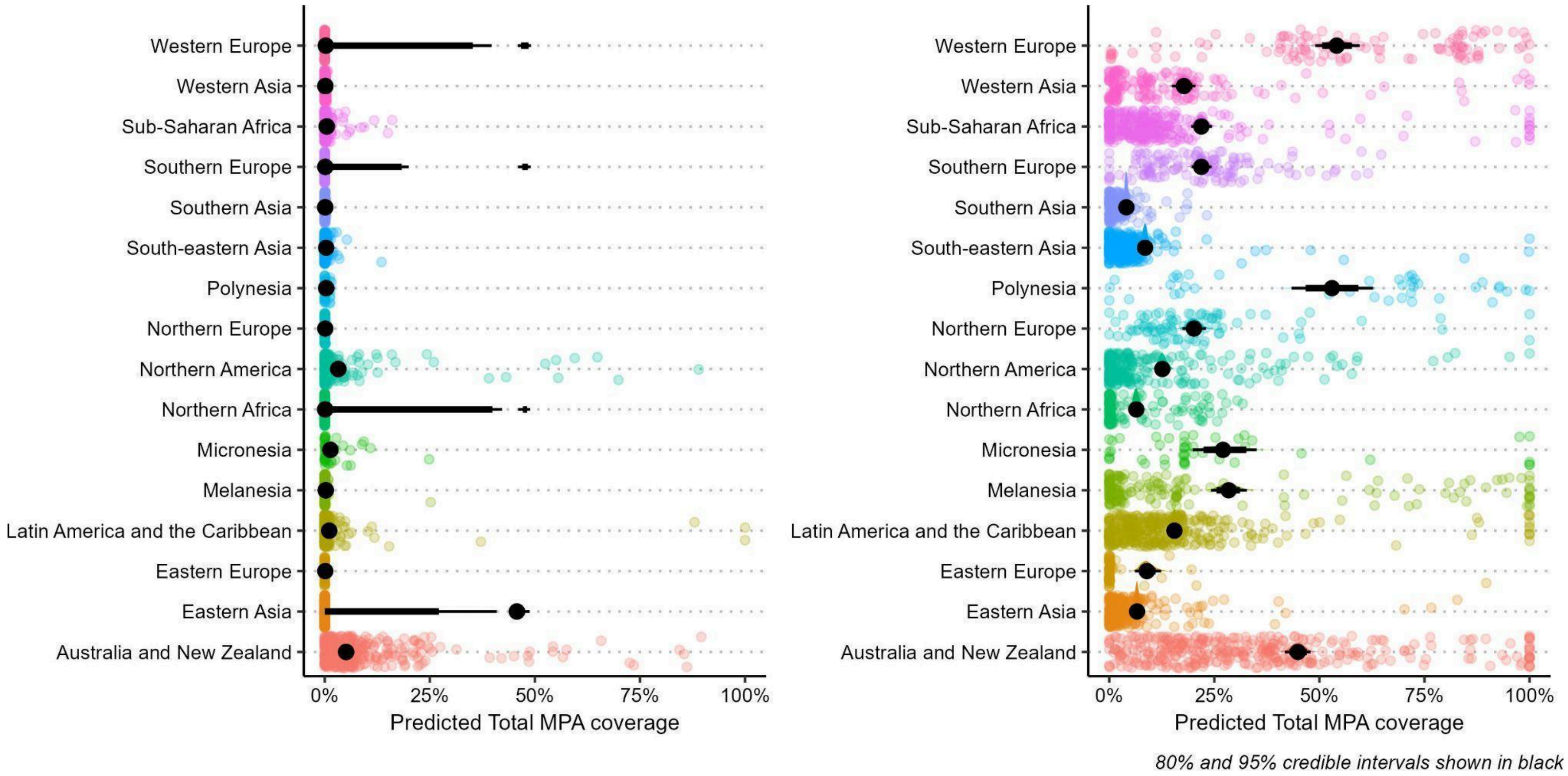


**Figure B6:** Relationship between subregion and predicted marine protected area (MPA) coverage for sharks and rays within associated EEZs. Predicted proportional coverage is shown for species within EEZs belonging to subregion categories. Points represent observed species-level values (n = 1,158). Black points indicate the posterior median predicted coverage from Bayesian zero–one inflated beta models, with black intervals showing the 80% and 95% credible intervals. Colored densities illustrate the posterior distribution of predicted values for each size category. The left panel shows predicted coverage within no-take MPAs only, whereas the right panel shows predicted total MPA coverage across all MPA categories.

**Appendix C: Data and Code Accessibility**

GitHub Repository

A public GitHub repository containing all code, including data processing, analyses, and plot generation, will be made available upon publication. The repository will include full documentation and instructions for replicating all results presented in the manuscript.

**Figure S1:** Screen capture of example query from SharkRayMPAexplorer Shiny App: What is the MPA coverage of each no-take category for threatened species in New Zealand?

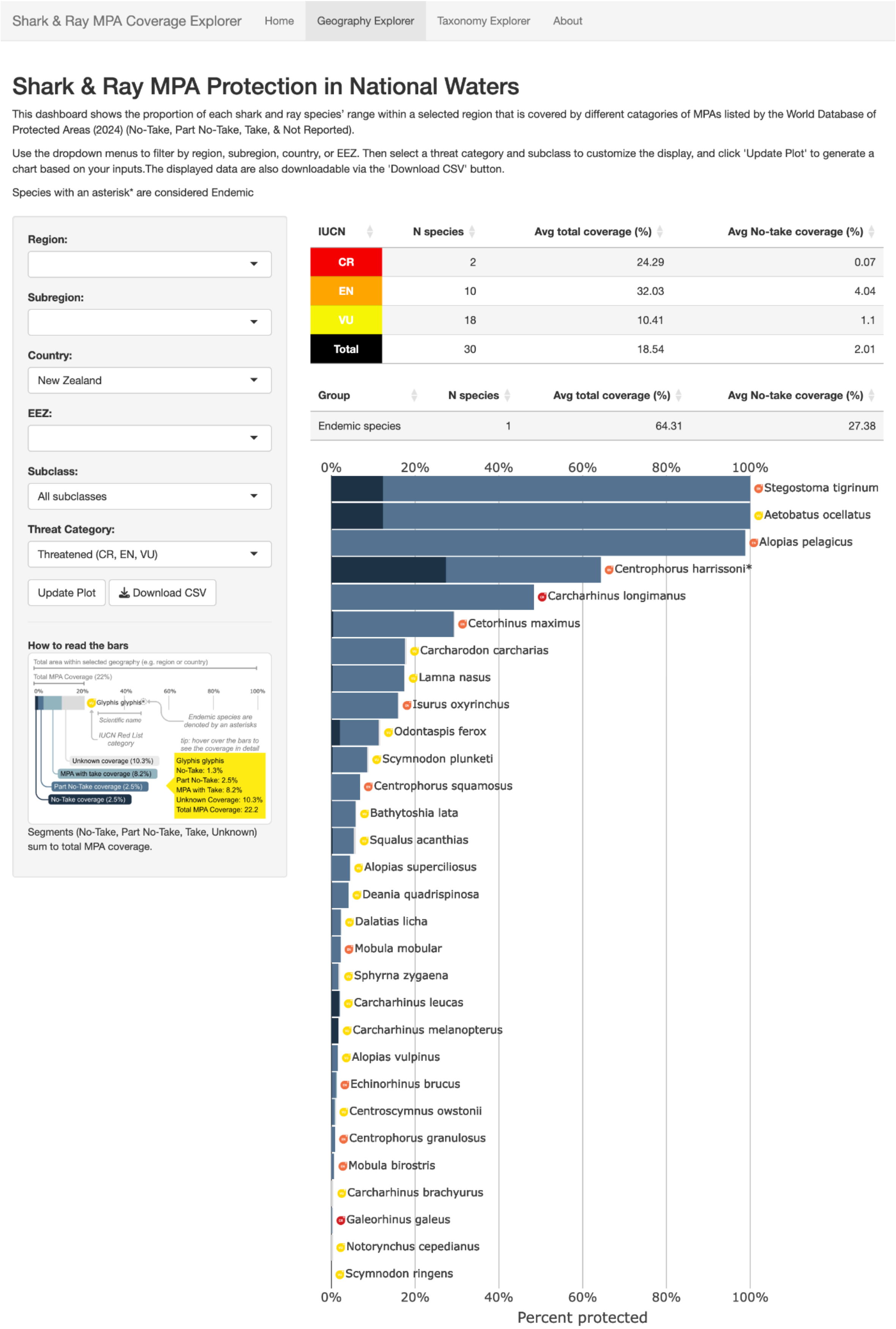

**Figure S2:** Screen capture of example query from SharkRayMPAexplorer Shiny App: What is the average no-take percent coverage for Rhinobatidae within global national waters?

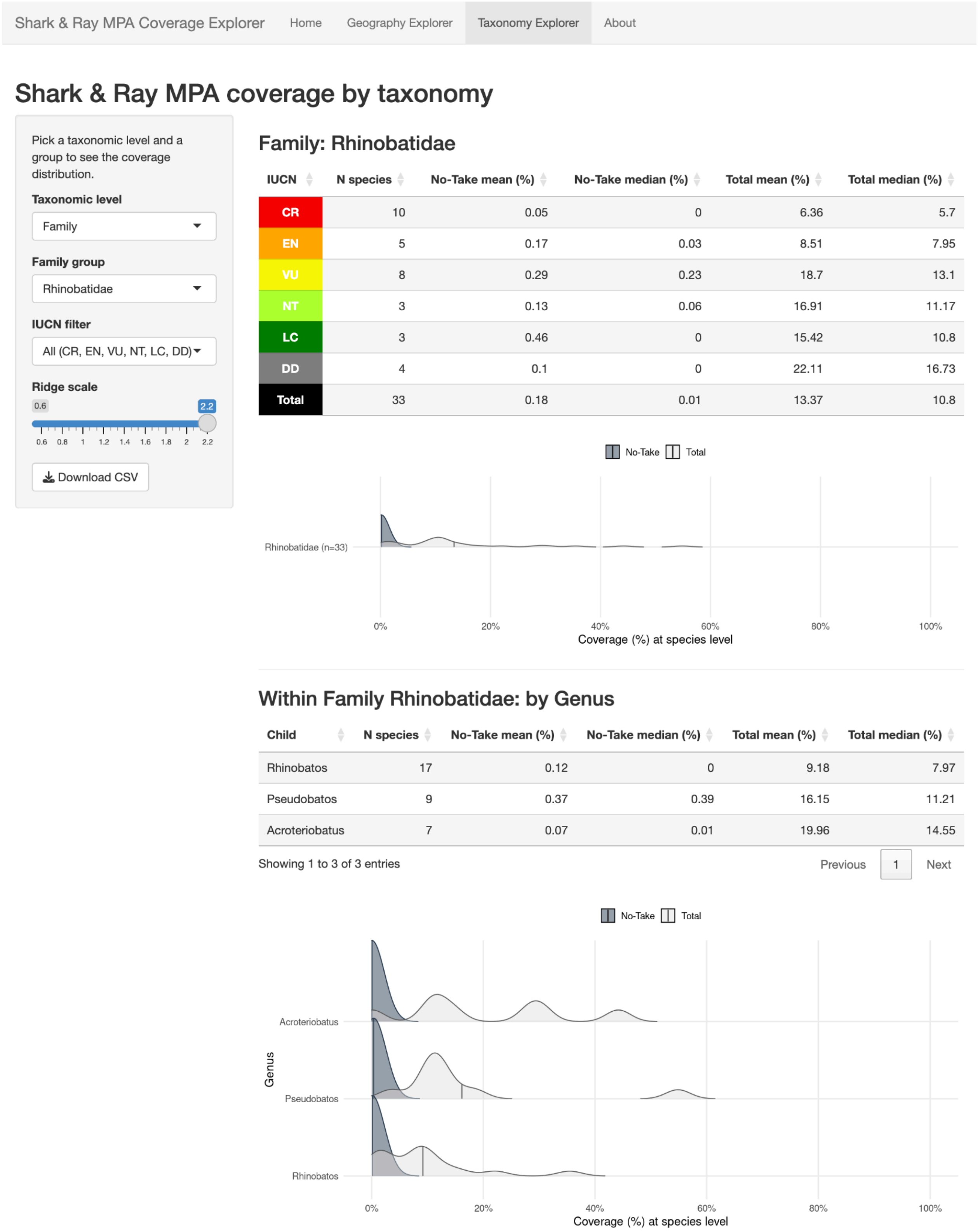



| IUCN | N species | No-Take mean (%) | No-Take median (%) | Total mean (%) | Total median (%) |
|---|---|---|---|---|---|
| CR | 10 | 0.05 | 0 | 6.36 | 5.7 |
| EN | 5 | 0.17 | 0.03 | 8.51 | 7.95 |
| VU | 8 | 0.29 | 0.23 | 18.7 | 13.1 |
| NT | 3 | 0.13 | 0.06 | 16.91 | 11.17 |
| LC | 3 | 0.46 | 0 | 15.42 | 10.8 |
| DD | 4 | 0.1 | 0 | 22.11 | 16.73 |
| Total | 33 | 0.18 | 0.01 | 13.37 | 10.8 |



| Child | N species | No-Take mean (%) | No-Take median (%) | Total mean (%) | Total median (%) |
|---|---|---|---|---|---|
| Rhinobatos | 17 | 0.12 | 0 | 9.18 | 7.97 |
| Pseudobatos | 9 | 0.37 | 0.39 | 16.15 | 11.21 |
| Acroteriobatus | 7 | 0.07 | 0.01 | 19.96 | 14.55 |

**Figure S3:** Map of countries with Marine Protected Areas (MPAs) listed in the WDPA that include a reported take status. Countries shaded in blue have varying levels of reporting, with color intensity scaled using a square-root transformation to highlight differences among countries with lower reporting levels. Countries shown in red have all MPAs classified as “Not Reported,” and countries in grey have no MPAs recorded in the WDPA.

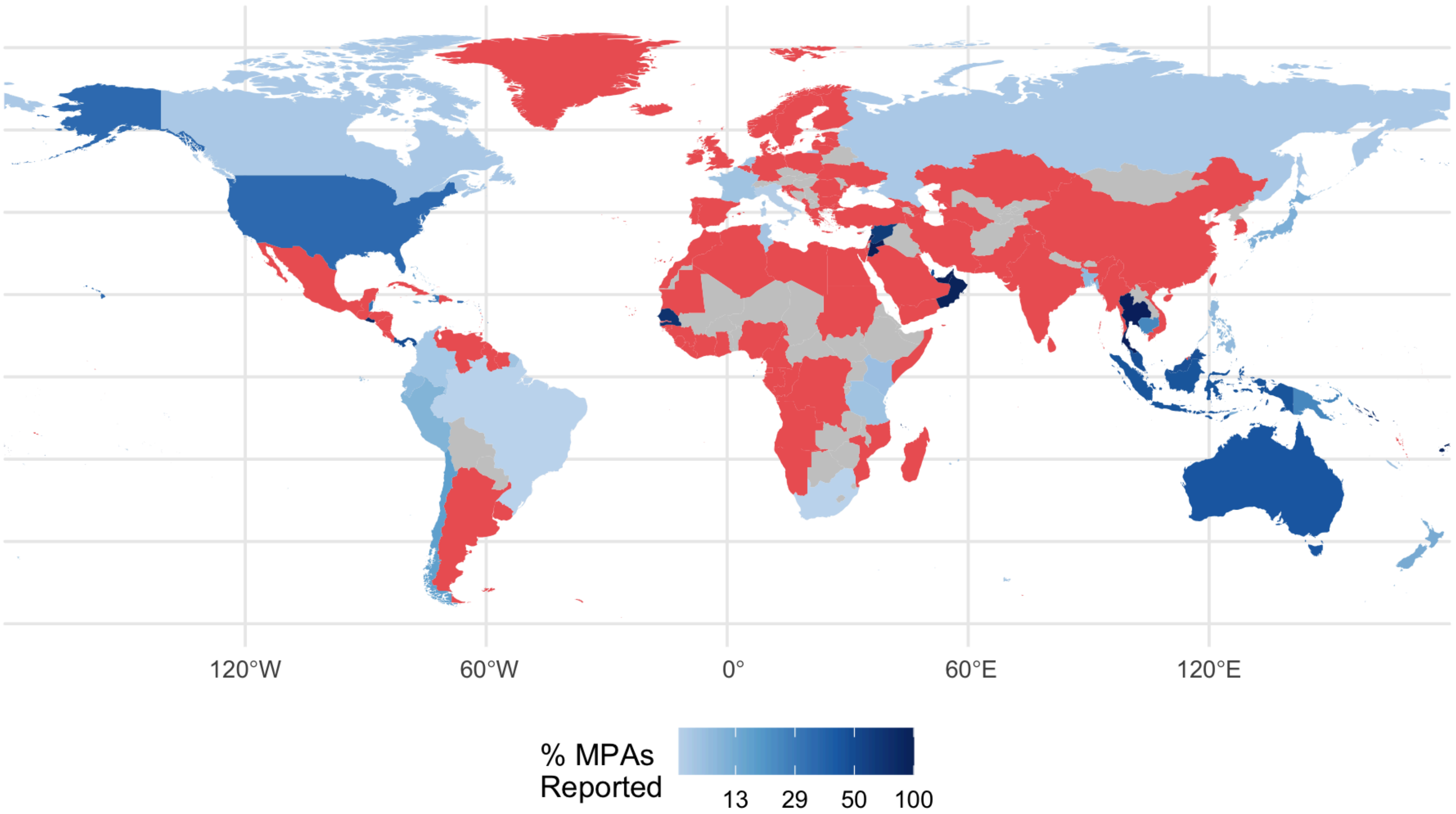

**Appendix D**: List of gap species.

Here, we separate no-take gap species and total gap species, and list all species with either 0% coverage or greater than 0 but less than 1% coverage.

<u>No-take gap species</u>

**Species with 0% no-take coverage (n=424):** Acroteriobatus blochii, Acroteriobatus salalah, Aculeola nigra, Aetobatus narutobiei, Amblyraja frerichsi, Amblyraja jenseni, Amblyraja reversa, Amblyraja taaf, Anacanthobatis marmorata, Apristurus aphyodes, Apristurus breviventralis, Apristurus canutus, Apristurus fedorovi, Apristurus gibbosus, Apristurus indicus, Apristurus investigatoris, Apristurus japonicus, Apristurus laurussonii, Apristurus macrorhynchus, Apristurus macrostomus, Apristurus manis, Apristurus microps, Apristurus nakayai, Apristurus ovicorrugatus, Apristurus profundorum, Apristurus saldanha, Apristurus stenseni, Apristurus yangi, Arhynchobatis asperrimus, Atelomycterus baliensis, Atelomycterus erdmanni, Atlantoraja castelnaui, Atlantoraja cyclophora, Atlantoraja platana, Bathyraja aguja, Bathyraja andriashevi, Bathyraja bergi, Bathyraja diplotaenia, Bathyraja fedorovi, Bathyraja hesperafricana, Bathyraja isotrachys, Bathyraja leucomelanos, Bathyraja macloviana, Bathyraja magellanica, Bathyraja matsubarai, Bathyraja pacifica, Bathyraja pallida, Bathyraja papilionifera, Bathyraja scaphiops, Bathyraja schroederi, Bathyraja smirnovi, Bathyraja smithii, Bathyraja trachouros, Bathyraja tzinovskii, Benthobatis kreffti, Benthobatis marcida, Benthobatis moresbyi, Beringraja pulchra, Breviraja claramaculata, Breviraja spinosa, Brevitrygon javaensis, Brochiraja aenigma, Brochiraja albilabiata, Brochiraja heuresa, Brochiraja microspinifera, Brochiraja vittacauda, Bythaelurus clevai, Bythaelurus giddingsi, Bythaelurus hispidus, Bythaelurus immaculatus, Bythaelurus lutarius, Bythaelurus tenuicephalus, Caliraja cortezensis, Carcharhinus borneensis, Centrophorus isodon, Centrophorus lesliei, Centrophorus longipinnis, Centroscyllium granulatum, Centroscyllium ornatum, Centroscyllium ritteri, Cephaloscyllium cooki, Cephaloscyllium fasciatum, Cephaloscyllium pictum, Cephaloscyllium sarawakensis, Cephaloscyllium silasi, Cephaloscyllium stevensi, Cephaloscyllium umbratile, Cephalurus cephalus, Chiloscyllium burmensis, Chiloscyllium caeruleopunctatum, Chimaera argiloba, Chimaera bahamaensis, Chimaera jordani, Chimaera notafricana, Chimaera opalescens, Chimaera owstoni, Chimaera supapae, Chimaera willwatchi, Chlamydoselachus africana, Cirrhigaleus barbifer, Cirrhoscyllium expolitum, Cirrhoscyllium japonicum, Cruriraja andamanica, Cruriraja atlantis, Cruriraja durbanensis, Cruriraja parcomaculata, Cruriraja poeyi, Ctenacis fehlmanni, Dasyatis hypostigma, Deania profundorum, Dichichthys albimarginatus, Dichichthys bigus, Dichichthys melanobranchus, Dichichthys nigripalatum, Dichichthys satoi, Diplobatis picta, Dipturus argentinensis, Dipturus campbelli, Dipturus chinensis, Dipturus crosnieri, Dipturus gigas, Dipturus johannisdavisi, Dipturus lamillai, Dipturus lanceorostratus, Dipturus leptocaudus, Dipturus macrocauda, Dipturus mennii, Dipturus nidarosiensis, Dipturus olseni, Dipturus oregoni, Dipturus oxyrinchus, Dipturus stenorhynchus, Dipturus tengu, Discopyge castelloi, Electrolux addisoni, Eridacnis radcliffei, Etmopterus alphus, Etmopterus benchleyi, Etmopterus brosei, Etmopterus burgessi, Etmopterus carteri, Etmopterus compagnoi,

Etmopterus decacuspidatus, Etmopterus lii, Etmopterus marshae, Etmopterus perryi, Etmopterus polli, Etmopterus robinsi, Etmopterus samadiae, Etmopterus sculptus, Etmopterus sentosus, Etmopterus splendidus, Euprotomicroides zantedeschia, Fenestraja atripinna, Fenestraja cubensis, Fenestraja ishiyamai, Fenestraja maceachrani, Fenestraja mamillidens, Fenestraja plutonia, Fontitrygon colarensis, Fontitrygon geijskesi, Fontitrygon ukpam, Galeus atlanticus, Galeus corriganae, Galeus eastmani, Galeus longirostris, Galeus mincaronei, Galeus murinus, Galeus nipponensis, Galeus piperatus, Galeus polli, Galeus schultzi, Glaucostegus granulatus, Glyphis gangeticus, Gurgesiella dorsalifera, Gurgesiella furvescens, Gymnura japonica, Gymnura micrura, Gymnura tentaculata, Halaelurus boesemani, Halaelurus buergeri, Halaelurus lineatus, Halaelurus maculosus, Harriotta raleighana, Hemiscyllium freycineti, Hemiscyllium galei, Hemiscyllium hallstromi, Hemiscyllium halmahera, Hemiscyllium henryi, Hemiscyllium michaeli, Hemitriakis complicofasciata, Hemitriakis indroyonoi, Hemitriakis japanica, Hemitriakis leucoperiptera, Hemitrygon akajei, Hemitrygon izuensis, Hemitrygon laevigata, Hemitrygon navarrae, Hemitrygon sinensis, Heterodontus japonicus, Heterodontus omanensis, Heterodontus quoyi, Heterodontus ramalheira, Heteronarce bentuviai, Heteronarce garmani, Heteronarce mollis, Holohalaelurus favus, Holohalaelurus grennian, Holohalaelurus melanostigma, Holohalaelurus punctatus, Hongeo koreana, Hydrolagus africanus, Hydrolagus alphus, Hydrolagus barbouri, Hydrolagus erithacus, Hydrolagus lusitanicus, Hydrolagus matallanasi, Hydrolagus mccoskeri, Hydrolagus mirabilis, Hydrolagus mitsukurii, Hydrolagus pallidus, Hydrolagus purpurescens, Hypanus berthalutzae, Hypanus marianae, Iago omanensis, Indobatis ori, Isogomphodon oxyrhynchus, Leucoraja circularis, Leucoraja compagnoi, Leucoraja erinacea, Leucoraja melitensis, Leucoraja yucatanensis, Maculabatis arabica, Maculabatis bineeshi, Malacoraja kreffti, Malacoraja obscura, Mollisquama mississippiensis, Mustelus albipinnis, Mustelus fasciatus, Mustelus griseus, Mustelus manazo, Mustelus minicanis, Mustelus schmitti, Mustelus sinusmexicanus, Mustelus whitneyi, Mustelus widodoi, Myliobatis chilensis, Myliobatis peruvianus, Myliobatis ridens, Myliobatis tobijei, Narcine baliensis, Narcine brasiliensis, Narcine insolita, Narcine oculifera, Narcine rierai, Narcine vermiculata, Narke japonica, Neoharriotta pinnata, Neoharriotta pumila, Neoraja africana, Neoraja caerulea, Neoraja carolinensis, Neoraja iberica, Neoraja stehmanni, Neotrygon yakkoei, Notoraja fijiensis, Notoraja hesperindica, Notoraja hirticauda, Notoraja inusitata, Notoraja longiventralis, Notoraja martinezi, Notoraja sereti, Notoraja sticta, Notoraja tobitukai, Okamejei acutispina, Okamejei boesemani, Okamejei cairae, Okamejei heemstrai, Okamejei hollandi, Okamejei kenojei, Okamejei meerdervoortii, Okamejei ornata, Okamejei picta, Okamejei schmidti, Orbiraja jensenae, Orbiraja powelli, Orectolobus japonicus, Oxynotus japonicus, Oxynotus paradoxus, Paragaleus leucolomatus, Parascyllium sparsimaculatum, Paratrygon orinocensis, Parmaturus albipenis, Parmaturus angelae, Parmaturus lanatus, Parmaturus pilosus, Pavoraja umbrosa, Pentanchus profundicolus, Planonasus indicus, Planonasus parini, Platyrhina hyugaensis, Platyrhina psomadakisi, Platyrhina sinensis, Platyrhina tangi, Pliotrema annae, Pristiophorus japonicus, Pristiophorus lanae, Pristiophorus nancyae, Pristiophorus schroederi, Proscyllium habereri, Proscyllium magnificum, Psammobatis bergi, Psammobatis extenta,

Psammobatis lentiginosa, Psammobatis normani, Psammobatis rutrum, Pseudobatos buthi, Pseudobatos horkelii, Raja herwigi, Raja maderensis, Raja polystigma, Rajella annandalei, Rajella bathyphila, Rajella bigelowi, Rajella dissimilis, Rajella eisenhardti, Rajella kukujevi, Rajella lintea, Rajella paucispinosa, Rajella purpuriventralis, Rajella ravidula, Rajella sadowskii, Rhinobatos annandalei, Rhinobatos austini, Rhinobatos holcorhynchus, Rhinobatos hynnicephalus, Rhinobatos jimbaranensis, Rhinobatos manai, Rhinobatos penggali, Rhinobatos ranongensis, Rhinobatos schlegelii, Rhinobatos whitei, Rhinochimaera atlantica, Rhinoraja kujiensis, Rhinoraja longicauda, Rhinoraja odai, Rhynchobatus cooki, Rhynchobatus immaculatus, Rhynchobatus mononoke, Rhynchorhina mauritaniensis, Rioraja agassizii, Rostroraja ackleyi, Rostroraja bahamensis, Schroederichthys chilensis, Schroederichthys saurisqualus, Schroederichthys tenuis, Scyliorhinus cabofriensis, Scyliorhinus hachijoensis, Scyliorhinus haeckelii, Scyliorhinus torazame, Scyliorhinus ugoi, Scylliogaleus quecketti, Scymnodalatias garricki, Scymnodon ichiharai, Scymnodon ringens, Sinobatis andamanensis, Sinobatis borneensis, Sinobatis brevicauda, Sinobatis melanosoma, Somniosus rostratus, Springeria folirostris, Springeria longirostris, Squalus acutipinnis, Squalus albicaudus, Squalus bahiensis, Squalus brevirostris, Squalus formosus, Squalus hawaiiensis, Squalus hemipinnis, Squalus japonicus, Squalus lalannei, Squalus lobularis, Squalus mahia, Squalus melanurus, Squalus mitsukurii, Squalus quasimodo, Squalus rancureli, Squalus shiraii, Squatina argentina, Squatina guggenheim, Squatina japonica, Squatina leae, Squatina mapama, Squatina nebulosa, Squatina occulta, Squatina tergocellatoides, Squatina varii, Sympterygia acuta, Sympterygia brevicaudata, Sympterygia lima, Telatrygon acutirostra, Telatrygon crozieri, Telatrygon zugei, Tetronarce formosa, Tetronarce puelcha, Tetronarce tokionis, Torpedo adenensis, Triakis acutipinna, Triakis maculata, Triakis scyllium, Urobatis concentricus, Urobatis maculatus, Urobatis pardalis, Urogymnus dalyensis, Urolophus aurantiacus, Urolophus deforgesi, Urolophus mitosis, Urolophus papilio, Urolophus sufflavus, Urotrygon cimar, Urotrygon microphthalmum, Zanobatus maculatus, Zapteryx brevirostris, Zearaja brevicaudata

**Species with <=1% no-take coverage (n=465):** Acroteriobatus annulatus, Acroteriobatus leucospilus, Acroteriobatus ocellatus, Acroteriobatus omanensis, Acroteriobatus variegatus, Acroteriobatus zanzibarensis, Aetobatus flagellum, Aetobatus laticeps, Aetobatus narinari, Aetomylaeus bovinus, Aetomylaeus maculatus, Aetomylaeus milvus, Aetomylaeus nichofii, Alopias superciliosus, Amblyraja doellojuradoi, Amblyraja georgiana, Amblyraja hyperborea, Amblyraja radiata, Apristurus brunneus, Apristurus exsanguis, Apristurus kampae, Apristurus longicephalus, Apristurus manocheriani, Apristurus melanoasper, Apristurus nasutus, Apristurus parvipinnis, Apristurus pinguis, Apristurus riveri, Apristurus sinensis, Asymbolus analis, Atelomycterus fasciatus, Atelomycterus marmoratus, Bathyraja abyssicola, Bathyraja albomaculata, Bathyraja aleutica, Bathyraja brachyurops, Bathyraja cousseauae, Bathyraja griseocauda, Bathyraja interrupta, Bathyraja kincaidii, Bathyraja lindbergi, Bathyraja maculata, Bathyraja meridionalis, Bathyraja microtrachys, Bathyraja minispinosa, Bathyraja multispinis, Bathyraja parmifera, Bathyraja peruana, Bathyraja richardsoni, Bathyraja shuntovi, Bathyraja spinicauda, Bathyraja taranetzi, Bathyraja trachura, Bathyraja violacea, Bathytoshia centroura, Beringraja binoculata, Breviraja colesi, Breviraja mouldi, Brevitrygon heterura, Brevitrygon manjajiae, Brevitrygon walga, Brochiraja asperula, Brochiraja leviveneta, Brochiraja spinifera, Bythaelurus canescens, Bythaelurus dawsoni, Caliraja inornata, Caliraja rhina, Caliraja stellulata, Callorhinchus callorynchus, Callorhinchus capensis, Carcharhinus acronotus, Carcharhinus brachyurus, Carcharhinus cerdale, Carcharhinus dussumieri, Carcharhinus humani, Carcharhinus isodon, Carcharhinus leiodon, Carcharhinus perezi, Carcharhinus porosus, Carcharhinus sealei, Carcharhinus signatus, Carcharhinus sorrah, Carcharhinus tjutjot, Centrophorus atromarginatus, Centrophorus seychellorum, Centrophorus squamosus, Centrophorus uyato, Centrophorus westraliensis, Centroscyllium fabricii, Centroscyllium nigrum, Centroscymnus coelolepis, Centroselachus crepidater, Cephaloscyllium isabellum, Cephaloscyllium sufflans, Cephaloscyllium ventriosum, Chaenogaleus macrostoma, Chiloscyllium arabicum, Chiloscyllium griseum, Chiloscyllium indicum, Chiloscyllium plagiosum, Chimaera carophila, Chimaera cubana, Chimaera fulva, Chimaera lignaria, Chimaera monstrosa, Chimaera ogilbyi, Chimaera orientalis, Chimaera phantasma, Chlamydoselachus anguineus, Cirrhigaleus asper, Cruriraja cadenati, Cruriraja hulleyi, Cruriraja rugosa, Dactylobatus clarkii, Dasyatis chrysonota, Dasyatis marmorata, Dasyatis pastinaca, Dasyatis tortonesei, Deania calcea, Deania quadrispinosa, Dentiraja australis, Dentiraja endeavouri, Dentiraja falloarga, Diplobatis ommata, Dipturus batis, Dipturus bullisi, Dipturus doutrei, Dipturus garricki, Dipturus grahamorum, Dipturus innominatus, Dipturus intermedius, Dipturus laevis, Dipturus pullopunctatus, Dipturus springeri, Dipturus teevani, Dipturus trachydermus, Discopyge tschudii, Echinorhinus brucus, Echinorhinus cookei, Eridacnis barbouri, Eridacnis sinuans, Etmopterus brachyurus, Etmopterus bullisi, Etmopterus fusus, Etmopterus gracilispinis, Etmopterus granulosus, Etmopterus hillianus, Etmopterus molleri, Etmopterus princeps, Etmopterus pusillus, Etmopterus schultzi, Etmopterus spinax, Etmopterus virens, Fenestraja sinusmexicanus, Fontitrygon margarita, Fontitrygon margaritella, Galeorhinus galeus, Galeus antillensis, Galeus arae, Galeus gracilis, Galeus melastomus, Galeus sauteri, Galeus springeri, Ginglymostoma cirratum, Ginglymostoma unami, Glaucostegus cemiculus, Glaucostegus halavi, Glaucostegus thouin, Glyphis glyphis, Gollum attenuatus, Gurgesiella atlantica, Gymnura altavela, Gymnura crebripunctata, Gymnura lessae, Gymnura marmorata, Gymnura natalensis, Gymnura poecilura, Gymnura sereti, Gymnura zonura, Halaelurus natalensis, Halaelurus quagga, Haploblepharus edwardsii, Haploblepharus fuscus, Haploblepharus kistnasamyi, Haploblepharus pictus, Harriotta avia, Harriotta haeckeli, Hemigaleus microstoma, Hemiscyllium strahani, Hemitriakis falcata, Hemitrygon bennettii, Hemitrygon longicauda, Hemitrygon parvonigra, Heterodontus francisci, Heterodontus marshallae, Heterodontus mexicanus, Heterodontus zebra, Hexanchus vitulus, Himantura leoparda, Himantura uarnak, Himantura undulata, Holohalaelurus regani, Hydrolagus affinis, Hydrolagus alberti, Hydrolagus bemisi, Hydrolagus colliei, Hydrolagus homonycteris, Hydrolagus macrophthalmus, Hydrolagus melanophasma, Hydrolagus novaezealandiae, Hypanus americanus, Hypanus dipterurus, Hypanus guttatus, Hypanus longus, Hypanus rudis, Hypanus sabinus, Hypanus say, Insentiraja subtilispinosa, Isistius brasiliensis, Isistius plutodus,

Lamiopsis temminckii, Lamiopsis tephrodes, Lamna ditropis, Leptocharias smithii, Leucoraja fullonica, Leucoraja garmani, Leucoraja leucosticta, Leucoraja naevus, Leucoraja pristispina, Leucoraja wallacei, Loxodon macrorhinus, Maculabatis ambigua, Maculabatis gerrardi, Maculabatis macrura, Maculabatis pastinacoides, Maculabatis randalli, Malacoraja spinacidermis, Megatrygon microps, Mitsukurina owstoni, Mobula hypostoma, Mobula kuhlii, Mobula mobular, Mustelus asterias, Mustelus californicus, Mustelus canis, Mustelus dorsalis, Mustelus henlei, Mustelus higmani, Mustelus lenticulatus, Mustelus lunulatus, Mustelus mento, Mustelus mosis, Mustelus mustelus, Mustelus norrisi, Mustelus palumbes, Mustelus punctulatus, Mustelus ravidus, Mustelus stevensi, Myliobatis aquila, Myliobatis californicus, Myliobatis freminvillei, Myliobatis goodei, Myliobatis longirostris, Narcine atzi, Narcine bancroftii, Narcine brevilabiata, Narcine entemedor, Narcine lingula, Narcine maculata, Narcine prodorsalis, Narcine timlei, Narcinops lasti, Narcinops ornata, Narke capensis, Narke dipterygia, Nasolamia velox, Negaprion brevirostris, Neotrygon annotata, Neotrygon caeruleopunctata, Neotrygon kuhlii, Neotrygon orientalis, Neotrygon varidens, Notoraja alisae, Notoraja azurea, Notorynchus cepedianus, Odontaspis ferox, Odontaspis noronhai, Okamejei leptoura, Orectolobus floridus, Orectolobus leptolineatus, Orectolobus reticulatus, Orectolobus wardi, Oxynotus caribbaeus, Oxynotus centrina, Paragaleus pectoralis, Paragaleus randalli, Paragaleus tengi, Parascyllium collare, Parmaturus macmillani, Parmaturus xaniurus, Pastinachus gracilicaudus, Pastinachus sephen, Pateobatis hortlei, Pateobatis jenkinsii, Pateobatis uarnacoides, Platyrhinoidis triseriata, Pliotrema warreni, Poroderma africanum, Poroderma pantherinum, Pristis clavata, Pristis pectinata, Pristis pristis, Psammobatis rudis, Psammobatis scobina, Pseudobatos glaucostigmus, Pseudobatos lentiginosus, Pseudobatos leucorhynchus, Pseudobatos percellens, Pseudobatos planiceps, Pseudobatos prahli, Pseudobatos productus, Pseudocarcharias kamoharai, Pseudoginglymostoma brevicaudatum, Pseudoraja fischeri, Pteroplatytrygon violacea, Raja asterias, Raja brachyura, Raja clavata, Raja microocellata, Raja miraletus, Raja montagui, Raja ocellifera, Raja parva, Raja radula, Raja straeleni, Raja undulata, Rajella barnardi, Rajella caudaspinosa, Rajella fuliginea, Rajella fyllae, Rajella leoparda, Rajella nigerrima, Rhinobatos albomaculatus, Rhinobatos borneensis, Rhinobatos irvinei, Rhinobatos lionotus, Rhinobatos punctifer, Rhinobatos rhinobatos, Rhinobatos sainsburyi, Rhinochimaera africana, Rhinochimaera pacifica, Rhinoptera bonasus, Rhinoptera brasiliensis, Rhinoptera javanica, Rhinoptera jayakari, Rhinoptera marginata, Rhinoptera steindachneri, Rhizoprionodon lalandii, Rhizoprionodon longurio, Rhizoprionodon oligolinx, Rhizoprionodon porosus, Rhizoprionodon terraenovae, Rhynchobatus djiddensis, Rhynchobatus laevis, Rhynchobatus luebberti, Rhynchobatus springeri, Rostroraja alba, Rostroraja cervigoni, Rostroraja eglanteria, Rostroraja equatorialis, Rostroraja texana, Rostroraja velezi, Schroederichthys bivius, Schroederobatis americana, Scoliodon laticaudus, Scoliodon macrorhynchos, Scyliorhinus boa, Scyliorhinus canicula, Scyliorhinus capensis, Scyliorhinus cervigoni, Scyliorhinus comoroensis, Scyliorhinus meadi, Scyliorhinus retifer, Scyliorhinus stellaris, Scyliorhinus torrei, Scymnodalatias albicauda, Scymnodalatias sherwoodi, Scymnodon macracanthus, Sinobatis bulbicauda, Sinobatis caerulea, Somniosus longus, Somniosus microcephalus, Sphyrna corona, Sphyrna lewini, Sphyrna media,

Sphyrna mokarran, Sphyrna tiburo, Sphyrna tudes, Sphyrna zygaena, Squaliolus laticaudus, Squalus acanthias, Squalus altipinnis, Squalus bassi, Squalus blainville, Squalus clarkae, Squalus crassispinus, Squalus cubensis, Squalus griffini, Squalus margaretsmithae, Squalus nasutus, Squalus suckleyi, Squatina aculeata, Squatina africana, Squatina armata, Squatina californica, Squatina david, Squatina dumeril, Squatina oculata, Squatina squatina, Styracura pacifica, Styracura schmardae, Sympterygia bonapartii, Taeniura lessoni, Taeniurops grabatus, Taeniurops meyeni, Telatrygon biasa, Tetronarce californica, Tetronarce cowleyi, Tetronarce occidentalis, Tetronarce tremens, Torpedo andersoni, Torpedo bauchotae, Torpedo fuscomaculata, Torpedo mackayana, Torpedo marmorata, Torpedo panthera, Torpedo sinuspersici, Torpedo torpedo, Triakis megalopterus, Triakis semifasciata, Trygonoptera imitata, Trygonorrhina fasciata, Typhlonarke aysoni, Urobatis halleri, Urobatis jamaicensis, Urogymnus acanthobothrium, Urogymnus lobistoma, Urogymnus polylepis, Urolophus kapalensis, Urolophus neocaledoniensis, Urotrygon aspidura, Urotrygon chilensis, Urotrygon munda, Urotrygon nana, Urotrygon reticulata, Urotrygon rogersi, Urotrygon simulatrix, Urotrygon venezuelae, Zameus squamulosus, Zanobatus schoenleinii, Zapteryx exasperata, Zapteryx xyster, Zearaja chilensis, Zearaja nasuta

**Species with 0% total coverage (n=74):** Amblyraja reversa, Apristurus investigatoris, Apristurus stenseni, Apristurus yangi, Bathyraja papilionifera, Bathyraja tzinovskii, Benthobatis kreffti, Breviraja claramaculata, Brochiraja albilabiata, Bythaelurus clevai, Bythaelurus immaculatus, Centroscyllium ornatum, Cephaloscyllium cooki, Cephaloscyllium stevensi, Chiloscyllium caeruleopunctatum, Chimaera bahamaensis, Chimaera supapae, Chimaera willwatchi, Cruriraja andamanica, Dichichthys nigripalatum, Dipturus johannisdavisi, Dipturus lamillai, Dipturus oregoni, Etmopterus decacuspidatus, Etmopterus lii, Etmopterus marshae, Etmopterus perryi, Etmopterus samadiae, Fenestraja maceachrani, Fenestraja mamillidens, Galeus corriganae, Galeus mincaronei, Gurgesiella dorsalifera, Halaelurus boesemani, Hemiscyllium halmahera, Hemiscyllium michaeli, Heteronarce mollis, Holohalaelurus melanostigma, Hydrolagus matallanasi, Malacoraja obscura, Mollisquama mississippiensis, Neoraja carolinensis, Notoraja hesperindica, Notoraja sereti, Okamejei picta, Orbiraja jensenae, Paratrygon orinocensis, Parmaturus angelae, Parmaturus lanatus, Pentanchus profundicolus, Planonasus indicus, Pristiophorus lanae, Pristiophorus schroederi, Rajella annandalei, Rajella paucispinosa, Rajella purpuriventralis, Rajella sadowskii, Rhinobatos manai, Rhynchobatus immaculatus, Rostroraja bahamensis, Sinobatis andamanensis, Sinobatis brevicauda, Springeria folirostris, Squalus bahiensis, Squalus hawaiiensis, Squalus lobularis, Squalus quasimodo, Squalus shiraii, Squatina leae, Squatina mapama, Squatina tergocellatoides, Torpedo adenensis, Urogymnus dalyensis, Urolophus mitosis

**Species with <=1% total coverage (n=69):** Acroteriobatus variegatus, Apristurus gibbosus, Apristurus indicus, Apristurus macrostomus, Apristurus profundorum, Bathyraja andriashevi, Bathyraja fedorovi, Bathyraja maculata, Bathyraja minispinosa, Bathyraja parmifera, Bathyraja scaphiops, Bathyraja smirnovi, Bathyraja taranetzi, Benthobatis moresbyi, Breviraja spinosa, Bythaelurus hispidus, Bythaelurus lutarius, Centrophorus isodon, Centrophorus longipinnis, Cephaloscyllium silasi, Chimaera argiloba, Chimaera jordani, Chimaera owstoni, Dichichthys melanobranchus, Dipturus chinensis, Dipturus crosnieri, Dipturus leptocaudus, Dipturus olseni, Etmopterus brosei, Etmopterus burgessi, Galeus eastmani, Glyphis glyphis, Halaelurus buergeri, Halaelurus quagga, Hemiscyllium hallstromi, Hemiscyllium strahani, Hemitrygon bennettii, Hemitrygon navarrae, Hongeo koreana, Hydrolagus lusitanicus, Hydrolagus purpurescens, Indobatis ori, Maculabatis arabica, Mustelus sinusmexicanus, Narcine brevilabiata, Narcine oculifera, Neoharriotta pumila, Neotrygon kuhlii, Notoraja inusitata, Okamejei acutispina, Okamejei boesemani, Parmaturus pilosus, Platyrhina psomadakisi, Platyrhina sinensis, Proscyllium habereri, Proscyllium magnificum, Rajella ravidula, Rhinobatos jimbaranensis, Rhinoraja kujiensis, Rhinoraja odai, Scyliorhinus ugoi, Scymnodon ichiharai, Sinobatis melanosoma, Squalus formosus, Squalus japonicus, Squalus rancureli, Squatina argentina, Taeniura lessoni, Urogymnus polylepis
